# Characterization of the workplace chemical exposome using untargeted LC-MS/MS: a case study

**DOI:** 10.1101/541813

**Authors:** Laura-Isobel McCall, Victoria M. Anderson, Robert S. Fogle III, Jacob J. Haffner, Ekram Hossain, Renmeng Liu, Anita H. Ly, Hongyan Ma, Maham Nadeem, Songyuan Yao

**Affiliations:** University of Oklahoma, Department of Chemistry and Biochemistry, 101 Stephenson Parkway, Norman, OK USA 73019; University of Oklahoma, Department of Microbiology and Plant Biology, 770 Van Vleet Oval, Norman, OK USA 73019; University of Oklahoma, Department of Anthropology, Dale Hall Tower Room 521, 455 West Lindsey, Norman, OK USA 73019; University of Oklahoma, Department of Cellular and Behavioral Neurobiology

**Keywords:** Built environment, Workplace chemical exposures, Untargeted metabolomics, Liquid chromatography-tandem mass spectrometry

## Abstract

Western people now spend close to 90% of their time indoors, one-quarter of which occurs at their place of employment. As such, interactions between employees and the workplace built environment have significant potential impact on employee health and safety. However, the range of workers’ daily chemical exposures is still poorly understood. Likewise, the influence of workers themselves and of worker behavior on the chemical composition of the workplace is still unknown. In this case study, we used untargeted liquid chromatography-tandem mass spectrometry (LC-MS/MS) to compare the chemical signatures of three different types of workplaces: scientific research buildings, office buildings, and one mixed-purpose building. Our results identified differential signatures of public building surfaces based on building purpose, sampling location and surface materials. Overall, these results are helping define the influence of human behavior on the workplace chemical environment and identify the chemical hazards to which people are exposed throughout their workday.

**Highlights:**

- Implementation of untargeted liquid chromatography-tandem mass spectrometry to study workplace chemical exposures.
- Shared chemical signatures were identified based on building purpose.
- Differential chemical signatures were identified based on surface material and sampling location.
- Annotated molecules include pharmaceuticals, illicit drugs, food chemicals, constituents of paints and stains, and cleaning products.

Abbreviations

LC-MS/MS: liquid chromatography-tandem mass spectrometry
*m/z*: mass over charge ratio
RT: retention time

## Introduction

Buildings are living spaces, housing vast numbers of bacteria in addition to their human occupants. There has been significant research into the microbiome of the built environment [1][2]. These studies showed distinct built environment microbiome based on room usage, such as for example distinctions between the microbiome of bathrooms, offices and kitchens [1][3][4], Human skin and outdoor environment are the major sources of the built environment microbiome [1]. However, these microbial surveys provide little insight into the functional consequences of these microbial colonizations. Metabolomic surveys of the built environment have the potential to identify not just products of microbial metabolism, but also the interactions between human building occupants and building surfaces. Such studies usually use mass spectrometry in combination with chromatographic separation (gas chromatography, or, increasingly, liquid chromatography), to identify and quantify the small molecules present in building air (e.g. [5]) or on building surfaces (*e.g.* [6][7][8]). In targeted metabolomics studies, researchers focus on a list of molecules of interest, usually known to be hazardous to human health. Such studies in the context of the built environment have for example quantified dust antimicrobial levels in houses and in workout rooms, hallways and offices of athletic facilities [9], or pesticide levels on household floors [6]. Untargeted metabolomics studies, in contrast, seek to detect the broadest possible range of molecules, with no *a priori* bias as to which molecules are interesting. Detected molecules include microbial products, but also compounds being leached by building surfaces (plasticizers, paint constituents…), cleaning products, and molecules deposited by building occupants themselves (*e.g.* beauty products, food derivatives) [7][8]. Such results can provide valuable information into a building’s usage and its occupants’ behavior.

While the majority of molecules found in buildings are likely innocuous, some can have an impact on people’s health. Workplace exposure to inhaled anesthetics for example is a known health risk for workers in the medical field [10], while dermal exposure to antimicrobials, detergents, dyes and disinfectants put healthcare and personal care workers at risk of occupational dermatitis [11]. Importantly, Petras et al revealed that laboratory chemicals are being spread outside of the laboratory [7], so that such molecules could have an impact not just on the health of the workers handling them, but also on visitors. We therefore sought to determine whether surface chemical risk exposures differ by building function. Surface samples were collected from public surfaces in two buildings dedicated to scientific research, two office buildings, and one mixed-purpose building housing teaching laboratories, offices and lecture rooms. Collected samples were analyzed by liquid chromatography-tandem mass spectrometry (LC-MS/MS) and grouped into chemical families using molecular networking. We observed that buildings with distinct purposes had different chemical profiles. Detected chemicals were also influenced by sampling location (floor vs door handles, for example) and surface material. Overall, these results illustrate the unique chemical risks to which building occupants are exposed depending on building purpose, and the interaction between building occupants and building surfaces. This data can help guide employee personal protection safeguards and inform building cleaning practices, while also providing insight into human behavior at sampled locations.

## Materials and Methods

### Sample collection

Two hundred and forty locations were sampled from five different buildings, including two laboratory, two office buildings and one high-traffic mixed-purpose building (housing offices, classrooms, and teaching laboratories). We refer to the laboratory buildings as buildings 1 and 2. Office buildings in this study are described as buildings 3 and 4, and the mixed-purpose building as building 5. The two laboratory and office buildings are within less than half a mile of each other within the same research park, and were all built as part of a concentrated construction effort. They are 2-14 years old. The mixed-purpose building is 2.3 miles away from the other buildings. It has been in constant use for the past 47 years The locations that were swabbed within each building were kept consistent and included: the right side of the main stair handrail going up, elevator buttons, the floor in front of six to nine labs or offices, the right armrest of couches, three wastebaskets, the outer door handle of three offices or labs, the inner building door handles, the floor by the exit door, light switches, and the water fountain. Each location swabbed was documented either with photos or detailed description. Cotton swabs were washed three times in 50% ethanol (all solvents were LC-MS grade) and soaked in 50% ethanol prior to use. The areas were swabbed for thirty seconds before placing the swabs in a deep 96-well plate containing 500 µL of 50% ethanol. For negative control, every twelfth sample was a blank swab in 50% ethanol. After samples had been collected, plates were sealed to prevent sample contamination and placed at 4oC overnight for further extraction. Swabs were then removed and extracts dried down (Thermo Fisher speedvac vacuum concentrator).

### Liquid chromatography- tandem mass spectrometry

LC-MS/MS sample preparation was performed by resuspending dried extracts in 50% methanol (spiked with 0.5 µg/mL sulfadimethoxine internal standard), with an injection volume of 20 µL. Column used in this analysis was a C18 core-shell column (Kinetex, 50×2.1 mm, 1.7 µM particle size, 100 Å pore size, Phenomenex, Torrance, USA). Mobile phase consisted of a two-solvent gradient (Solvent A: H_2_O+0.1% formic acid, Solvent B: Acetonitrile+0.1% formic acid). Gradient parameters were: 5% B for 1 min, then linear increase from 5% B to 100% B over 8 minutes, hold at 100% B for 2 minutes and return to 5% B in 30 seconds, with a subsequent 1 min re-equilibration phase at 5% B. Column temperature was maintained at 40°C and sample compartment at 10°C for the entirety of the analysis. Samples were run in randomized order with blanks every 12 samples; blanks alternated between swab blanks (blank swab extracted with 50% ethanol) and plate blanks (50% methanol plus internal standard only). Electrospray (ESI) parameters were set at 35 L/min, 10 L/min auxiliary gas flow rate, 0 L/min sweep gas flow rate, and 350°C auxiliary gas temperature. The spray voltage was set to 3.8 kV, S-lens RF level was at 50 V and the capillary temperature was set at 320°C. Data was acquired in positive mode, with data-dependent MS2 acquisition. The MS scans had a scan range of 100-1500 m/z and 5 MS/MS scans of the most abundant ion per cycle were recorded. Resolution for MS1 was set to 35,000 and 17,500 for MS2. Maximum injection time for both MS1 and MS2 was set at 100 ms. Full MS AGC target was 1e6. MS/MS AGC target was 5e5. An isolation window of 2 m/z was selected. Normalized collision energy was incrementally increased from 20% to 30% and to 40%. MS/MS occurred at an apex of 2-8 seconds with a dynamic exclusion of 10 seconds. Last, ions with unassigned charges were excluded from instrumental analysis.

### Data analysis

Raw MS data files were converted to mzXML format using MSconvert software (http://proteowizard.sourceforge.net/tools.shtml). MS features were identified using MZmine (v. 2.33) using parameters shown in Table 1 [12]. Only features with abundance >5 times abundance in blank swab samples were retained. Total ion current (TIC) normalization was performed using the R language implemented in Jupyter Notebook ((http://jupyter.org/)). Principal Coordinate Analysis (PCoA) plots of MS features were created from a Canberra dissimilarity matrix, using an in-house clustering script. Distance matrices were obtained using QIIME 1 [12,13], and PERMANOVA calculations performed using the R package “vegan”. To identify differential features between locations, the 1,000 most abundant features were examined using random forest machine learning approaches in R in Jupyter Notebooks, using 5,000 trees and classifying based on building type. Cross-validation was performed by splitting the data 80-20 using the R package “caret”, training the random forest model on 80% of the data (training dataset), and then assessing classification accuracy on the remainder of the dataset (test dataset). Data log-transformed using MetaboAnalyst (https://www.metaboanalyst.ca)[14] was analyzed by one-way or Welch’s ANOVA, depending on the within-group variance, using in-house developed R script. Molecular networking was performed using the Global Natural Products Social Molecular Networking (GNPS) online platform [15] on the .mgf file exported from MZmine, using the following parameters: precursor ion and fragment ion mass tolerance: 0.02 Da; minimum cosine score for networking and library matching: 0.7; minimum number of matched MS2 fragment ions for networking and library matching: 4; network topK: 50; maximum connected component size: 100; analog search: enabled; maximum analog mass difference: 100 Da; precursor window filtering: enabled; 50 Da peak window filtering: enabled; row sum normalization. Libraries searched were all GNPS spectral libraries, METLIN, LipidBlast, NIST_17, and an in-house library of contaminants. These search parameters are associated with less than 5% false discovery rates [16]; annotations were further curated manually based on mirror plot appearance and plausibility of the chemical changes for analog matches. Generated annotations are at level 2/3 as defined by the metabolomics standard initiative (putatively annotated compounds or compound classes) [17]. Networks were visualized using Cytoscape version 3.7.0 [18]. Matching to previous studies of the built environment or humans was performed using the single spectrum search option in GNPS, with the following search parameters: parent and fragment ion tolerance, 0.02 Da; minimum matched peaks: 4; score threshold: 0.7; do not search unclustered data or analogs; precursor and 50 Da peak window filtering, enabled. Chemical structures were generated using ChemDraw software (Perkin Elmer).

**Table 1.** MZmine Parameters used in Data Analysis.

|  |  |  |
| --- | --- | --- |
| Mass Detection | MS1 Noise Level | 1.20E+05 |
|  | MS2 Noise Level | 1E+03 |
|  | Mass Detector | Centroid |
| Chromatogram Builder<br>Baseline cutoff algorithm | Minimum Time Span (min) | 0.05 |
|  | Minimum Height | 1.70E+06 |
|  | m/z tolerance (ppm) | 10 |
| Chromatogram Deconvolution | Minimum Peak Height | 1.70E+06 |
|  | Peak Duration Range | 0.00-2.00 |
|  | Baseline Level | 1.00E+06 |
|  | m/z Range for MS2 Scan Pairing (Da) | 0.01 |
|  | RT Range for MS2 Scan Pairing (min.) | 0.10 |
| Isotopic Peak Grouper | Retention Time Tolerance (min) | 0.10 |
|  | *m/z* tolerance (ppm) | 10 |
|  | Monotonic Shape | Yes |
|  | Maximum Charge | 3 |
|  | Representative isotope | Lowest *m/z* |
| Join aligner | *m/z* tolerance (ppm) | 10 |
|  | *m/z* to RT weight | 5 to 1 |
|  | Retention Time Tolerance (min) | 0.10 |
| Row filtering | Retention Time | 0.20-12 min |
|  | Keep only peaks with MS2 scan | enabled |

### Data availability

LC-MS/MS data has been deposited in MassIVE under accession number MSV000082953. Molecular network can be accessed at https://gnps.ucsd.edu/ProteoSAFe/status.jsp?task=70c1775687724c8cacc0a324208a91c4 (overall analysis) and https://gnps.ucsd.edu/ProteoSAFe/status.jsp?task=540b7367604648b0941b6ac37ac5e314 (dataset matching).

## Results

### Surface metabolite profile segregates by building usage

To determine the impact of building use on a building’s chemical profile, we collected surface chemicals from five buildings, all within 2.3 miles of each other, including two buildings used for scientific research, two office buildings, and one mixed-purpose building housing classrooms, teaching laboratories and offices. Samples were collected by swabbing public areas of the buildings, including stair handrail, elevator buttons, floors outside offices or lab, couches, light switches, garbage can lids and door handles. Molecules were extracted from swabs using 50% ethanol and analyzed by liquid chromatography-tandem mass spectrometry, followed by processing using MZmine2 [12] to extract molecular features, and molecular networking for feature annotation and grouping into chemical families [15]. Overall, this analytical approach detected 23,030 molecular features, which were grouped into 2,599 chemical families and 8,736 singletons (features not grouped into families) (Fig. S6).

To evaluate and visualize the similarities between samples from different building types, principal coordinate analysis (PCoA) was performed. PCoA analysis showed clustering based on the different building types (**Fig. 1A** PERMANOVA p<0.001, **Fig. S3**), indicating common chemical profiles based on building usage and a clear distinction between research, mixed-purpose and office buildings. We further subset our dataset to only perform PCoA analysis on office *vs* research buildings. Results showed distinct clustering (indicating different overall chemical composition) between office and research buildings, with some of the most differential samples coming from door handles and stairway railings (**Fig. 1B** PERMANOVA p<0.001). In accordance with our hypothesis of shaping of the building surface metabolome by building function, there was considerable overlap between our mixed-purpose building and other building types, as evidenced by close clustering of samples from the mixed-purpose building with samples from the other building types by PCoA analysis (**Fig. 1A**), higher mis-classification of building 5-derived samples by our random forest classifier (**Fig. 1C**), and lower frequency of molecular features unique to this building (**Fig. 1G, Fig. S1**). Indeed, 11.33% of molecular features identified in our mixed-purpose building were also identified in research buildings, and 6.05% were shared between the mixed-purpose building and office buildings.

**Figure 1.**
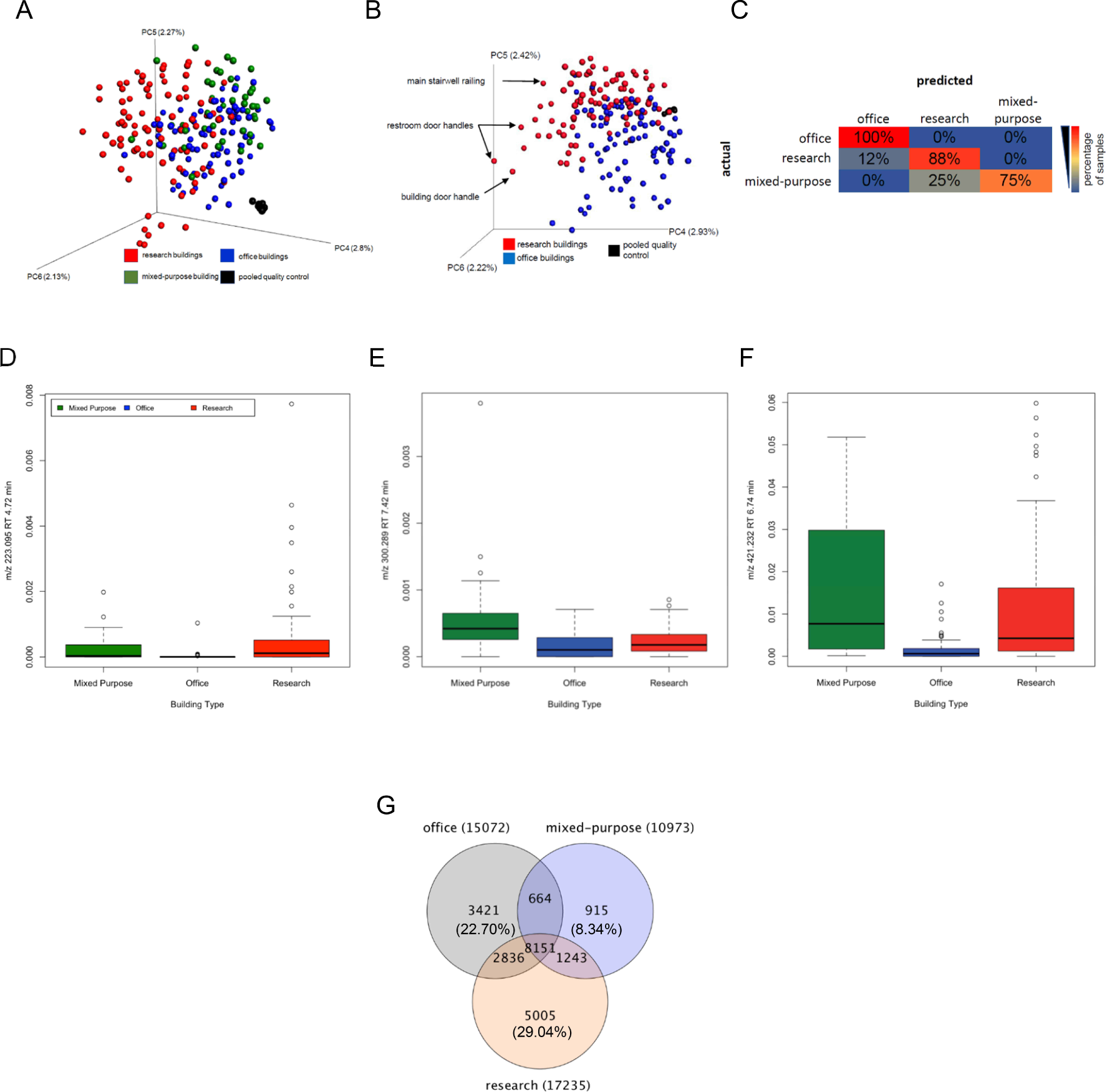
Differential building surface chemical profile depending on building function. (A) Principal coordinate analysis showing partial segregation of samples based on building type, comparing office, research and mixed-purpose buildings (Canberra distance metric; p<0.001 PERMANOVA). (B) Principal coordinate analysis showing partial segregation of samples based on building type, comparing office and research buildings only (Canberra distance metric; p<0.001PERMANOVA). (C) Random Forest classification results on test dataset. High classification accuracy was obtained, indicating that all three building types present distinct chemical profiles. Correct classification is along the diagonal. Percentage of samples classified into each category are displayed. (D-F) Representative differentially-abundant molecules between building types, as identified by random forest analysis: diethyl phthalate (D), palmitoyl ethanolamide (E), tris(2-butoxyethyl) phosphate (F). (G) Venn diagram showing features unique to each building type, with the lowest proportion of unique molecules found in our mixed-purpose building.

Next, we sought to determine which molecular features were key “signatures” of building usage. We performed random forest classification analysis [19] on the top 1000 most abundant features in our dataset. Random forest analysis showed excellent classification accuracy (**Fig. 1C**), supporting the presence of differential chemical profiles based on building usage, in accordance with our PCoA analysis results. Most of the chemicals identified by random forest as differing between buildings based on building purpose could not be annotated, but many have been reported in other studies of the built environment (**Table 2, Fig. S2**). Strikingly, several molecular features, including *m/z* 272.258 RT 4.75 min, *m/z* 425.252 RT 3.91 min and *m/z* 819.474 RT 6.74 min, all of which were highest in the research and mixed-purpose buildings (**Fig. S4**), were previously detected on water fountains from a research building (MassIVE dataset MSV000079720, [7]). *m/z* 470.369 RT 6.12 was found across all building types in our study, with the highest levels in our office buildings (**Fig. S4**); in accordance with these observations, it was reported in a variety of built environment settings: research building water fountain, apartment and researcher’s office inside a science building (**Table 2**). Piperine, a food-derived molecule, was reported in human studies and in an analysis of apartment surfaces; we detected it in our mixed-purpose building. In contrast, several of our differential surface features have not yet been reported on studies of the built environment but have been detected in human-derived samples. While the lack of reports in the context of the built environment may merely reflect the bias of much of current metabolomics research towards human analysis, these shared reports suggest molecular exchange between building occupants and building surfaces. For example, *m/z* 279.232 RT 7.06, annotated as the plant-derived fatty acid linolenic acid and palmitoyl ethanolamide (*m/z* 300.289 RT 7.42 min), a human-produced fatty acid amide, were both highest in our high-traffic mixed-purpose building and previously identified in human metabolomics studies. In contrast, detection of chemicals such as tris(2-butoxyethyl) phosphate, a flame retardant and plasticizer, and diethyl phthalate (*m/z* 223.095 RT 4.72 min), a plasticizer, sealant and coating constituent, in prior human studies suggest transfer of chemicals from built environment surfaces to humans. Other notable differential molecules include a derivative of N-(2-Hydroxypropyl)dodecanamide (cosmetic constituent) and food derivatives at higher levels in mixed-purpose and research buildings. This may reflect building occupant behavior and the higher human traffic into these buildings compared to our office locations.

**Table 2.**
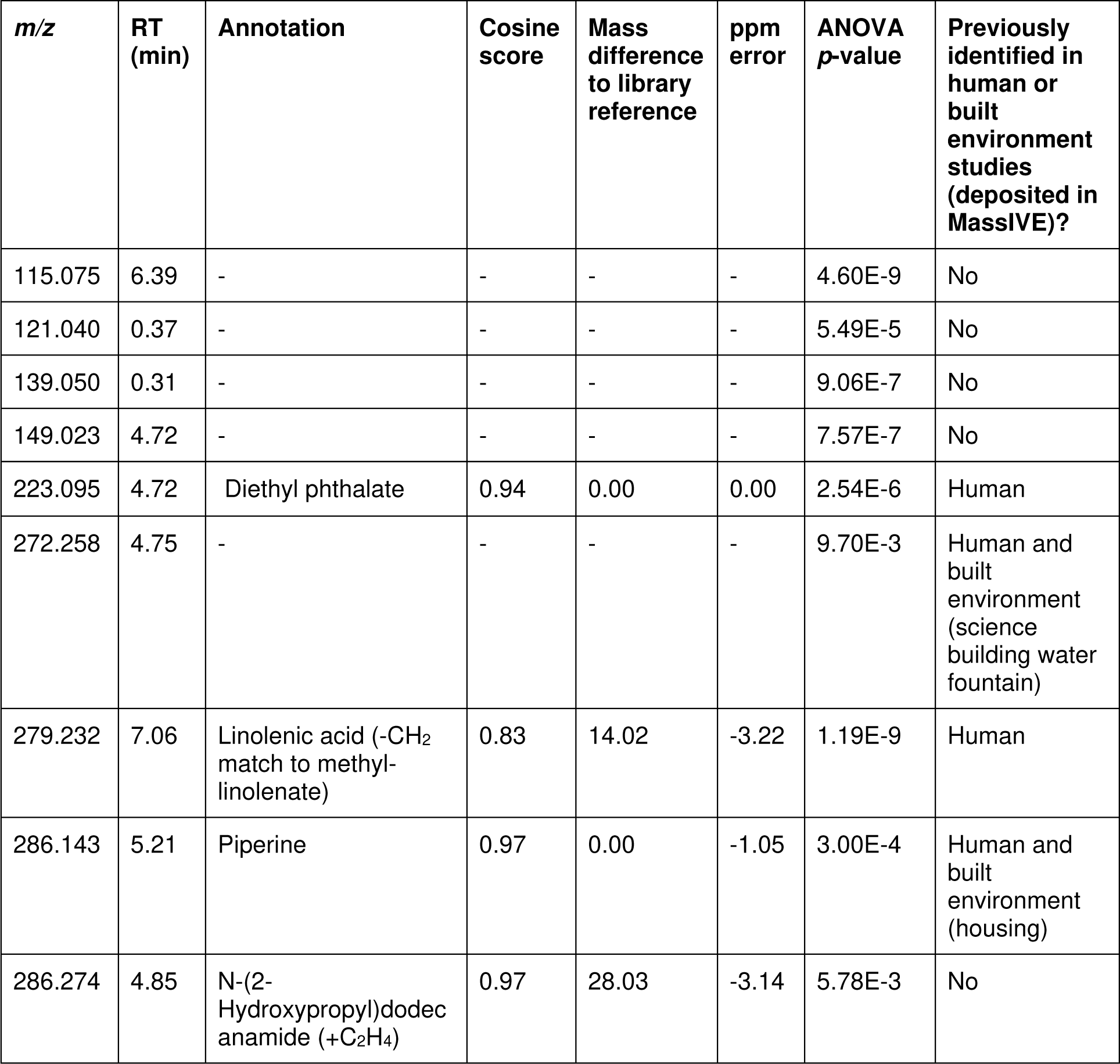
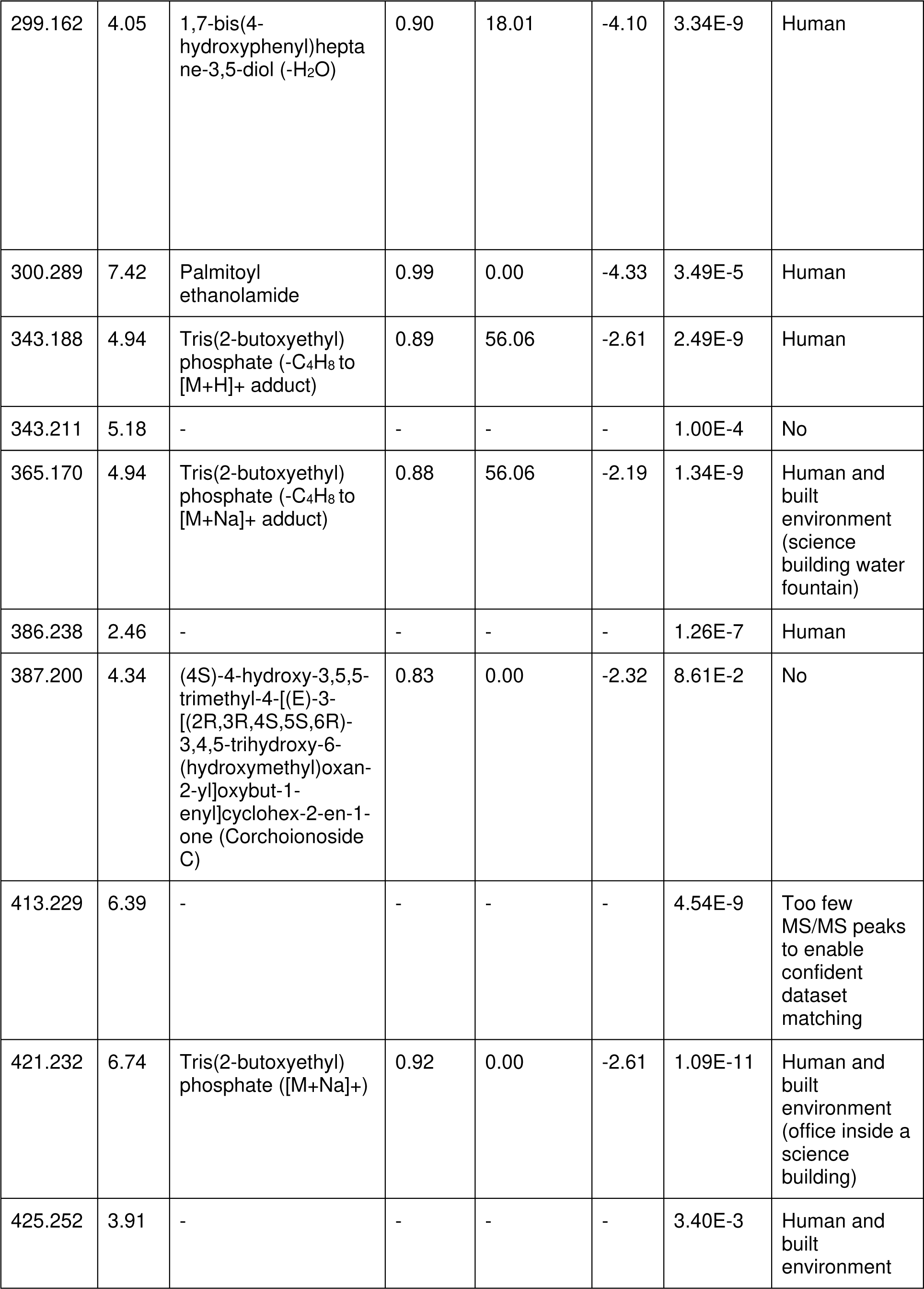
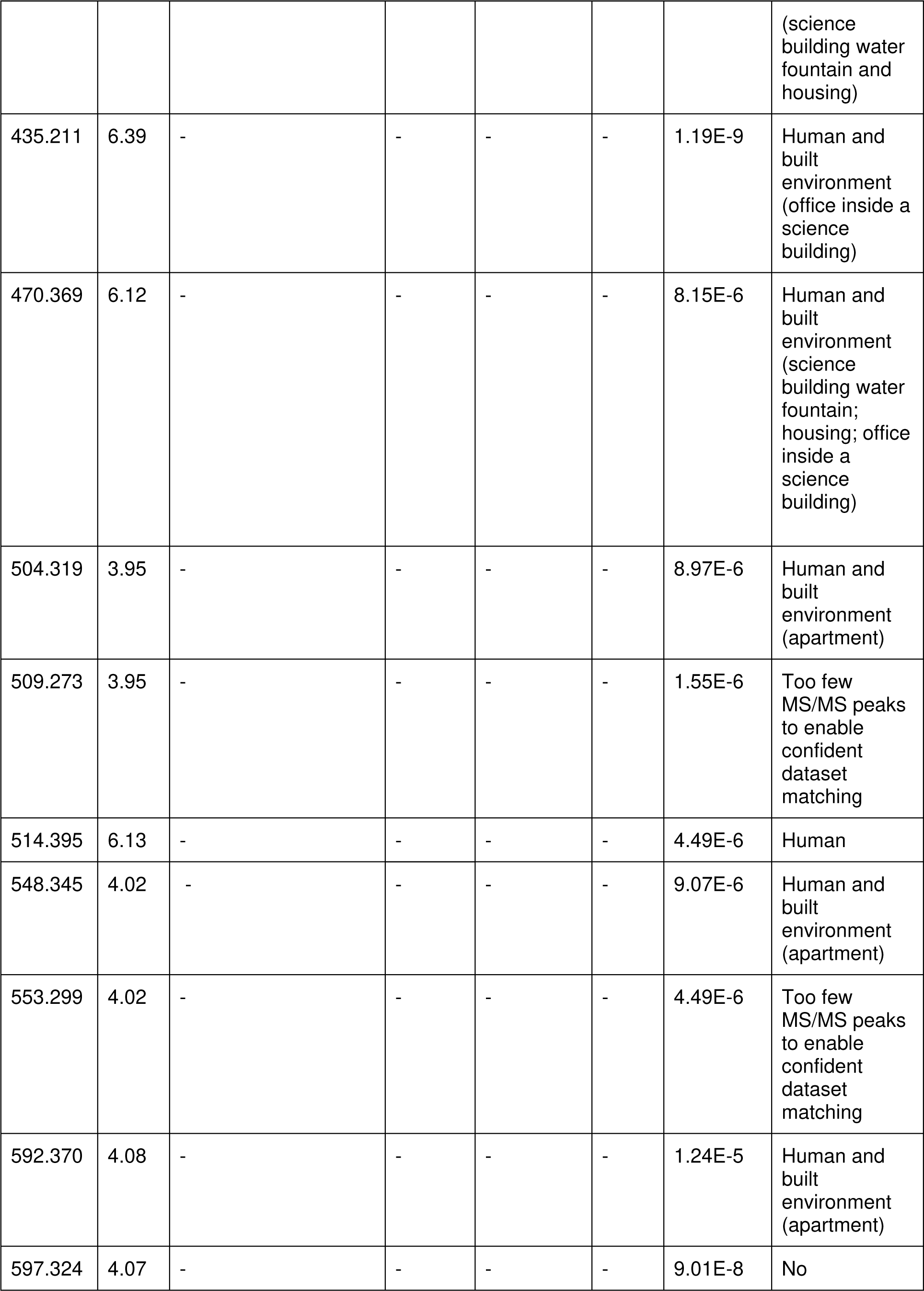
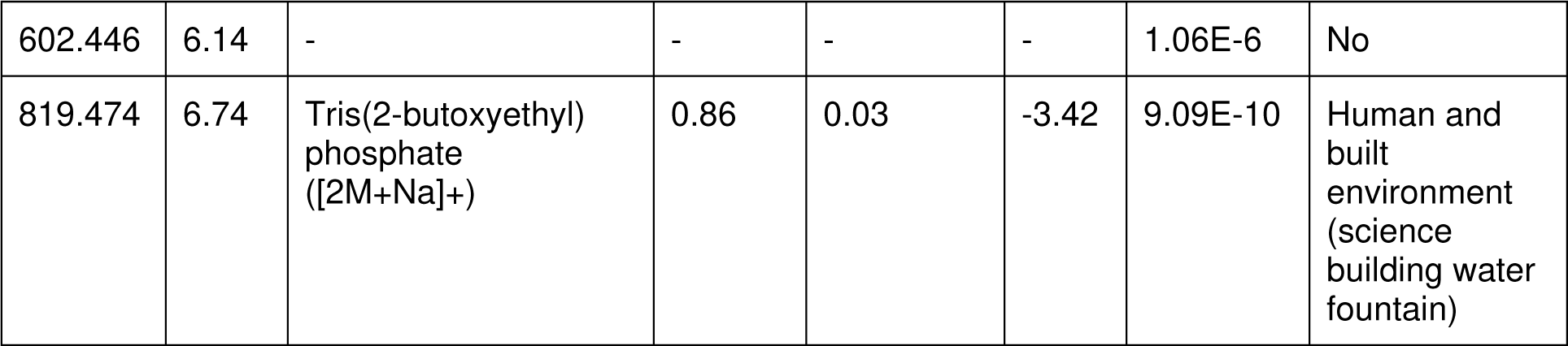
Top 30 most differential features, as identified by random forest classifier.

| *m/z* | RT (min) | Annotation | Cosine score | Mass difference to library reference | ppm error | ANOVA *p*-value | Previously identified in human or built environment studies (deposited in MassIVE)? |
| --- | --- | --- | --- | --- | --- | --- | --- |
| 115.075 | 6.39 | - | - | - | - | 4.60E-9 | No |
| 121.040 | 0.37 | - | - | - | - | 5.49E-5 | No |
| 139.050 | 0.31 | - | - | - | - | 9.06E-7 | No |
| 149.023 | 4.72 | - | - | - | - | 7.57E-7 | No |
| 223.095 | 4.72 | Diethyl phthalate | 0.94 | 0.00 | 0.00 | 2.54E-6 | Human |
| 272.258 | 4.75 | - | - | - | - | 9.70E-3 | Human and built environment (science building water fountain) |
| 279.232 | 7.06 | Linolenic acid (-CH<sub>2</sub> match to methyl-linolenate) | 0.83 | 14.02 | -3.22 | 1.19E-9 | Human |
| 286.143 | 5.21 | Piperine | 0.97 | 0.00 | -1.05 | 3.00E-4 | Human and built environment (housing) |
| 286.274 | 4.85 | N-(2-Hydroxypropyl)dodecanamide (+C<sub>2</sub>H<sub>4</sub>) | 0.97 | 28.03 | -3.14 | 5.78E-3 | No |
| 299.162 | 4.05 | 1,7-bis(4-hydroxyphenyl)heptane-3,5-diol (-H<sub>2</sub>O) | 0.90 | 18.01 | -4.10 | 3.34E-9 | Human |
| 300.289 | 7.42 | Palmitoyl ethanolamide | 0.99 | 0.00 | -4.33 | 3.49E-5 | Human |
| 343.188 | 4.94 | Tris(2-butoxyethyl) phosphate (-C<sub>4</sub>H<sub>8</sub> to [M+H]<sup>+</sup> adduct) | 0.89 | 56.06 | -2.61 | 2.49E-9 | Human |
| 343.211 | 5.18 | - | - | - | - | 1.00E-4 | No |
| 365.170 | 4.94 | Tris(2-butoxyethyl) phosphate (-C<sub>4</sub>H<sub>8</sub> to [M+Na]<sup>+</sup> adduct) | 0.88 | 56.06 | -2.19 | 1.34E-9 | Human and built environment (science building water fountain) |
| 386.238 | 2.46 | - | - | - | - | 1.26E-7 | Human |
| 387.200 | 4.34 | (4S)-4-hydroxy-3,5,5-trimethyl-4-[(E)-3-[(2R,3R,4S,5S,6R)-3,4,5-trihydroxy-6-(hydroxymethyl)oxan-2-yl]oxybut-1-enyl]cyclohex-2-en-1-one (Corchoionoside C) | 0.83 | 0.00 | -2.32 | 8.61E-2 | No |
| 413.229 | 6.39 | - | - | - | - | 4.54E-9 | Too few MS/MS peaks to enable confident dataset matching |
| 421.232 | 6.74 | Tris(2-butoxyethyl) phosphate ([M+Na]<sup>+</sup>) | 0.92 | 0.00 | -2.61 | 1.09E-11 | Human and built environment (office inside a science building) |
| 425.252 | 3.91 | - | - | - | - | 3.40E-3 | Human and built environment (science building water fountain and housing) |
| 435.211 | 6.39 | - | - | - | - | 1.19E-9 | Human and built environment (office inside a science building) |
| 470.369 | 6.12 | - | - | - | - | 8.15E-6 | Human and built environment (science building water fountain; housing; office inside a science building) |
| 504.319 | 3.95 | - | - | - | - | 8.97E-6 | Human and built environment (apartment) |
| 509.273 | 3.95 | - | - | - | - | 1.55E-6 | Too few MS/MS peaks to enable confident dataset matching |
| 514.395 | 6.13 | - | - | - | - | 4.49E-6 | Human |
| 548.345 | 4.02 | - | - | - | - | 9.07E-6 | Human and built environment (apartment) |
| 553.299 | 4.02 | - | - | - | - | 4.49E-6 | Too few MS/MS peaks to enable confident dataset matching |
| 592.370 | 4.08 | - | - | - | - | 1.24E-5 | Human and built environment (apartment) |
| 597.324 | 4.07 | - | - | - | - | 9.01E-8 | No |
| 602.446 | 6.14 | - | - | - | - | 1.06E-6 | No |
| 819.474 | 6.74 | Tris(2-butoxyethyl) phosphate ([2M+Na]⁺) | 0.86 | 0.03 | -3.42 | 9.09E-10 | Human and built environment (science building water fountain) |

Finally, we considered molecules unique to a given building. Such molecules can provide insight into the unique activities of that building’s occupants. For example, we found many pharmaceuticals and illicit drugs in our high-traffic building. These included erythromycin (antibacterial), cyclobenzaprine (muscle relaxant) on a building entrance door handle and cocaine in many sampling locations. Plant-derived molecules such as caryophyllene oxide or oleanolic acid were also found in several locations. Locations varied from floors to doors and also included elevator interiors. Pharmaceuticals and illicit drugs were most commonly found on high-touch surfaces (e.g. doorknobs/handles), however illicit drugs were also more prevalent on surfaces which did not receive as much cleaning (a wooden statue, for example). (**Table 3, Fig. S10**). Overall, these results provide a snapshot into the diversity of possible chemical exposures when entering any given building.

**Table 3.**
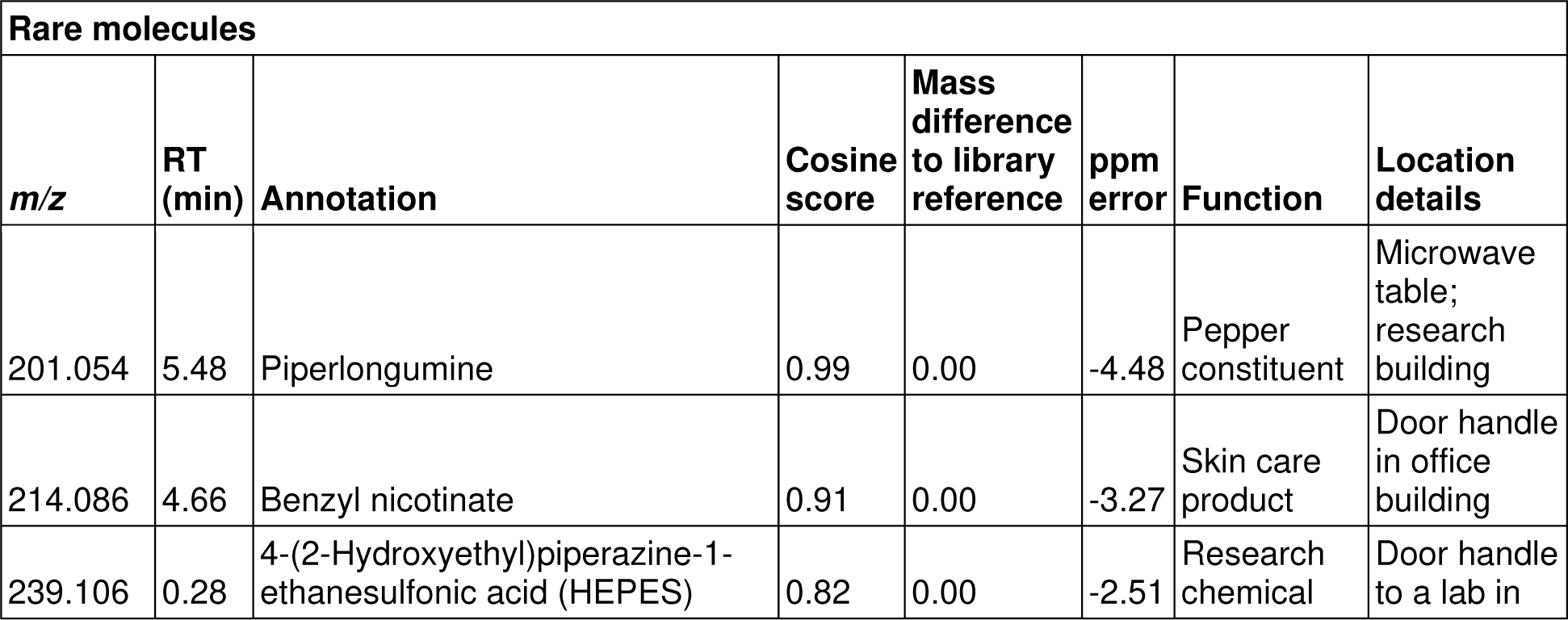
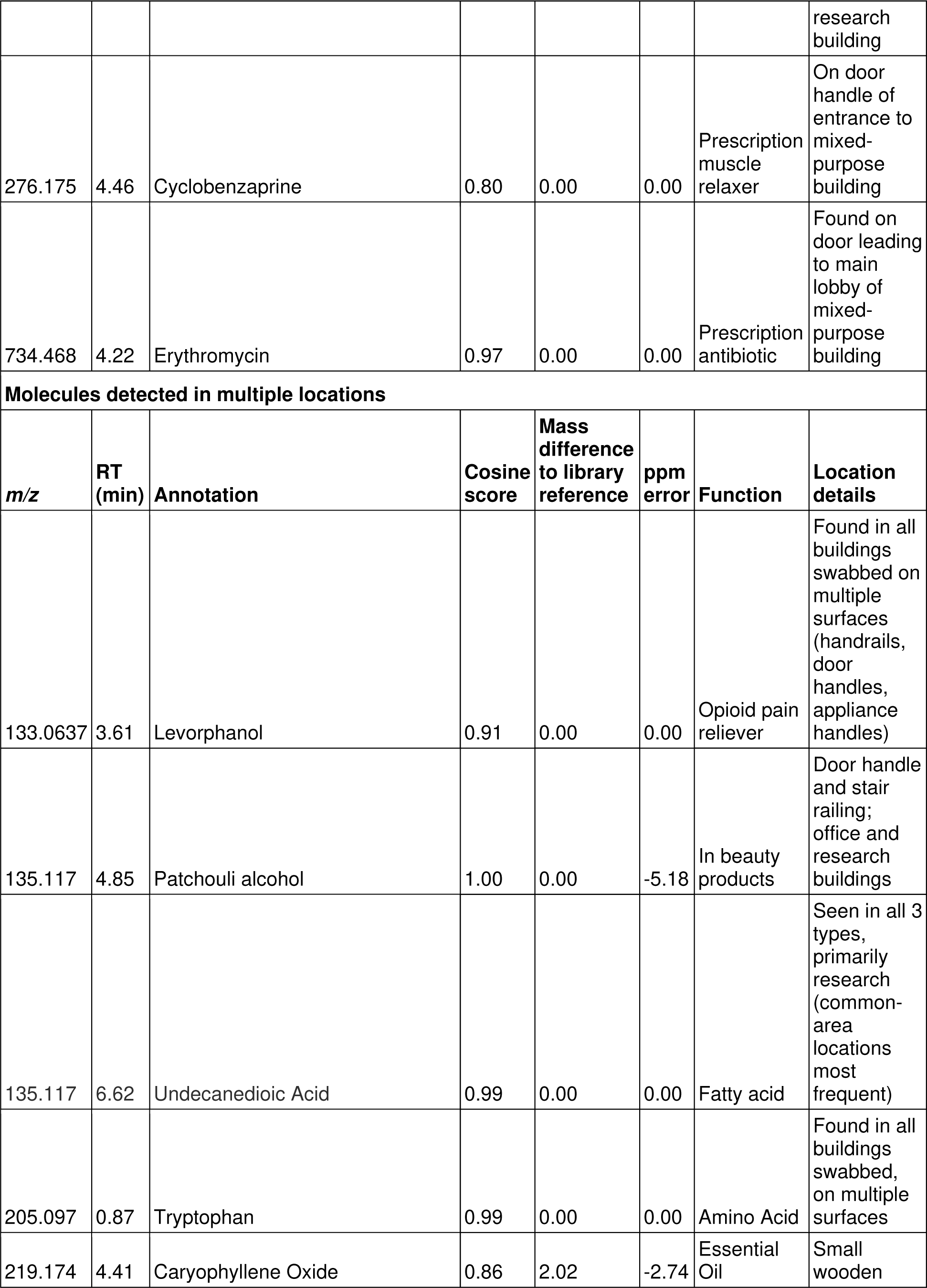
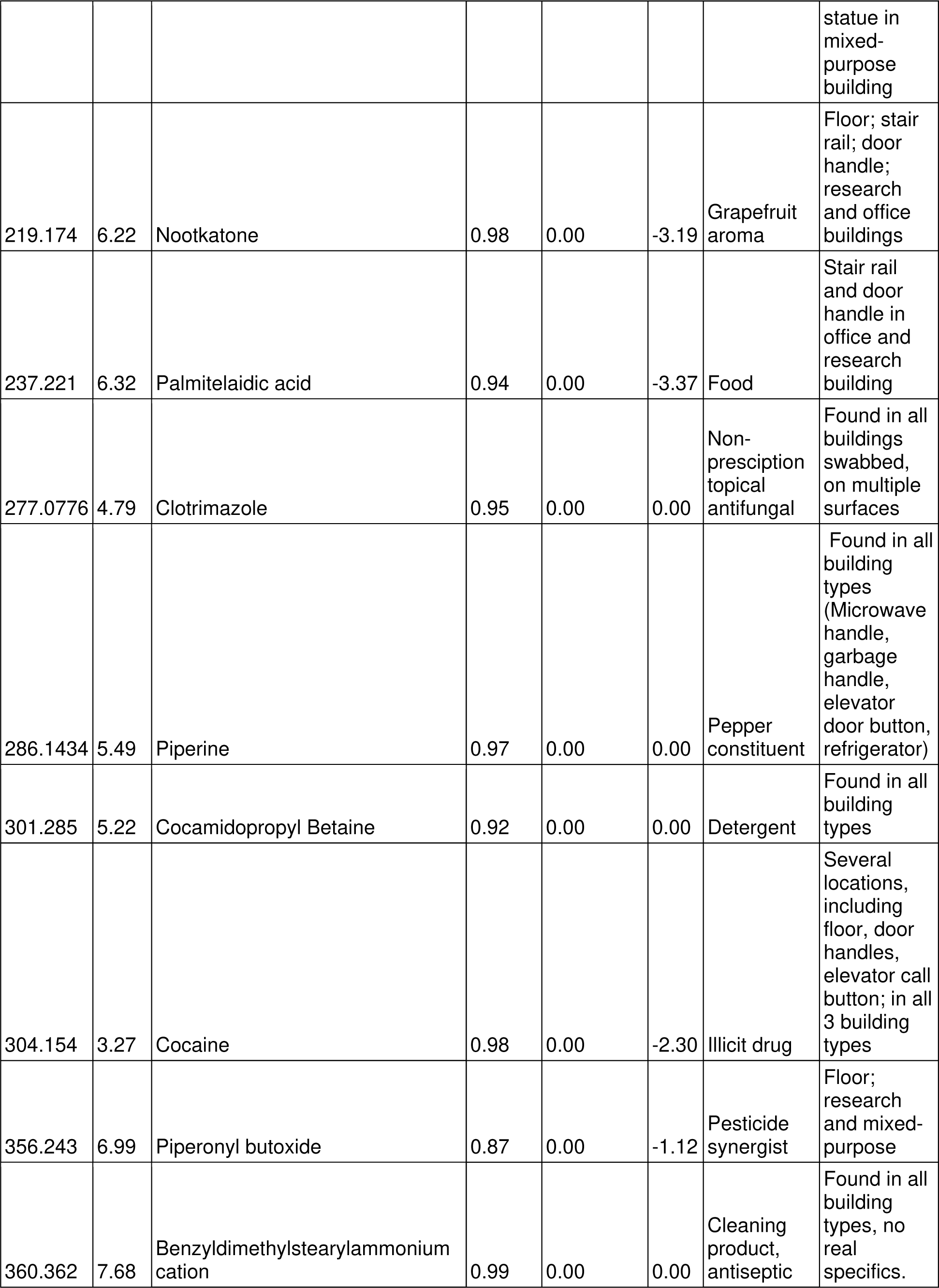
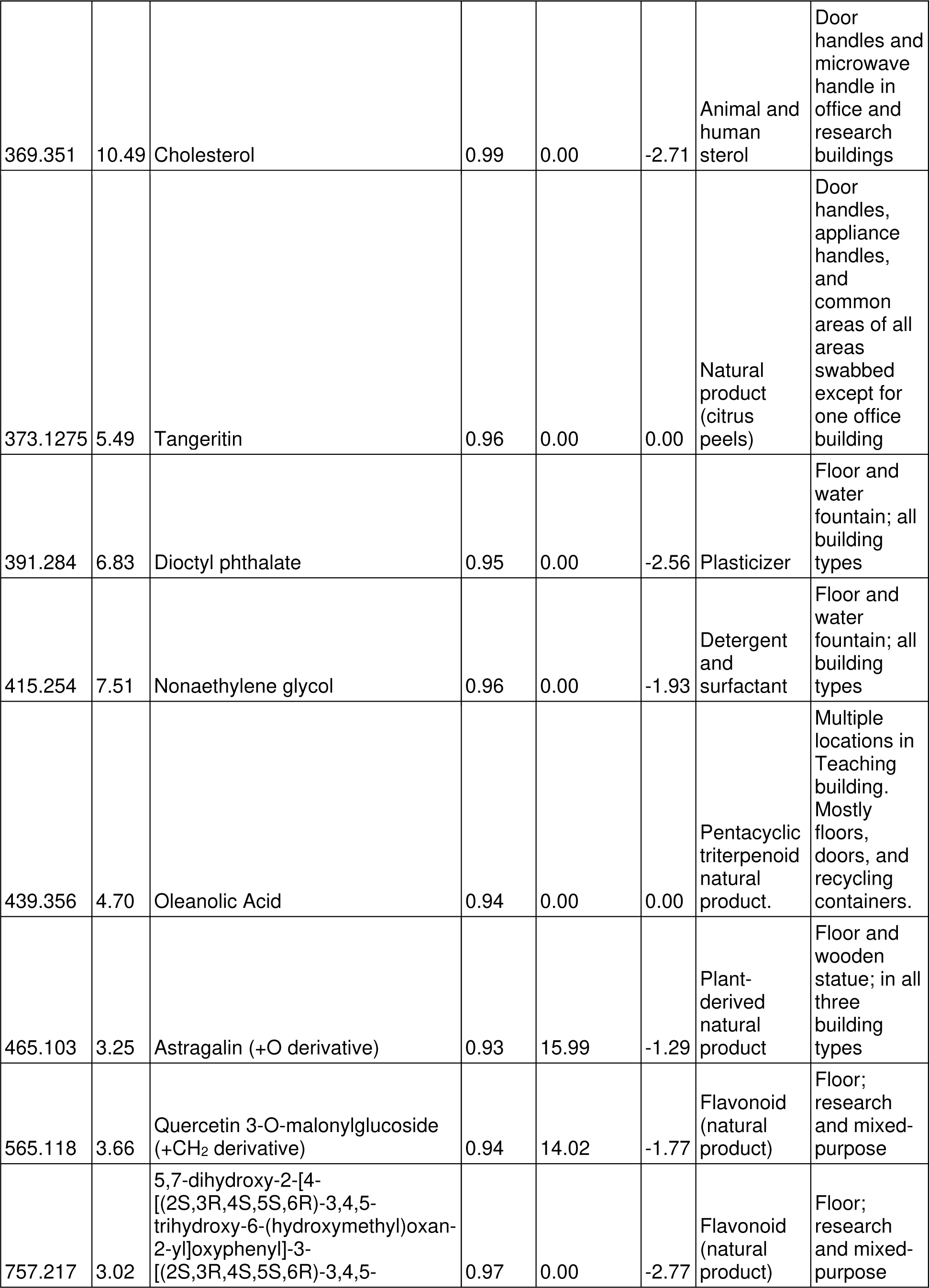
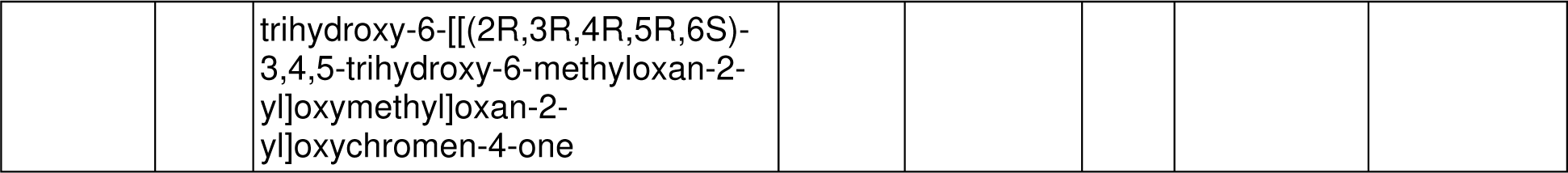
Additional molecules of interest detected on building surfaces.

### Common chemical signatures based on sampling location

Samples were collected from a variety of surfaces and locations within buildings. We also observed significant heterogeneity in recovered molecule abundance even within a given building (**Fig. S4**), suggesting an impact of the location sampled within a given building. We therefore visualized the impact of sampling location by PCoA analysis. Overall, we observed significant differences in overall chemical profile between surfaces on which people walk (floors in front of offices and labs, elevator floor) and surfaces people touch (handrails, door handles, elevator buttons etc.) (**Fig. 2A**, PERMANOVA p<0.001). Examples of molecules commonly found on floors but not on door handles include pesticide constituents (piperonyl butoxide, *m/z* 356.243 RT 6.99 min); detergents (nonaethylene glycol, *m/z* 415.254 RT 7.51 min) and plant-derived molecules (astragalin derivative, *m/z* 465.1027 RT 3.25 min; some flavonoids (*m/z* 565.118 RT 3.66 min and *m/z* 757.217 RT 3.02 min; **Table 3, Fig. S10**). Likewise, molecules found on surfaces people touch but not on floors include 4-(2-Hydroxyethyl)piperazine-1-ethanesulfonic acid (HEPES; research chemical found in laboratory building; *m/z* 239.106 RT 0.28 min); patchouli alcohol (*m/z* 135.117 RT 4.85 min) and cholesterol (*m/z* 369.351 RT 10.49 min) (**Table 3, Fig. Sxx**).

**Figure 2.**
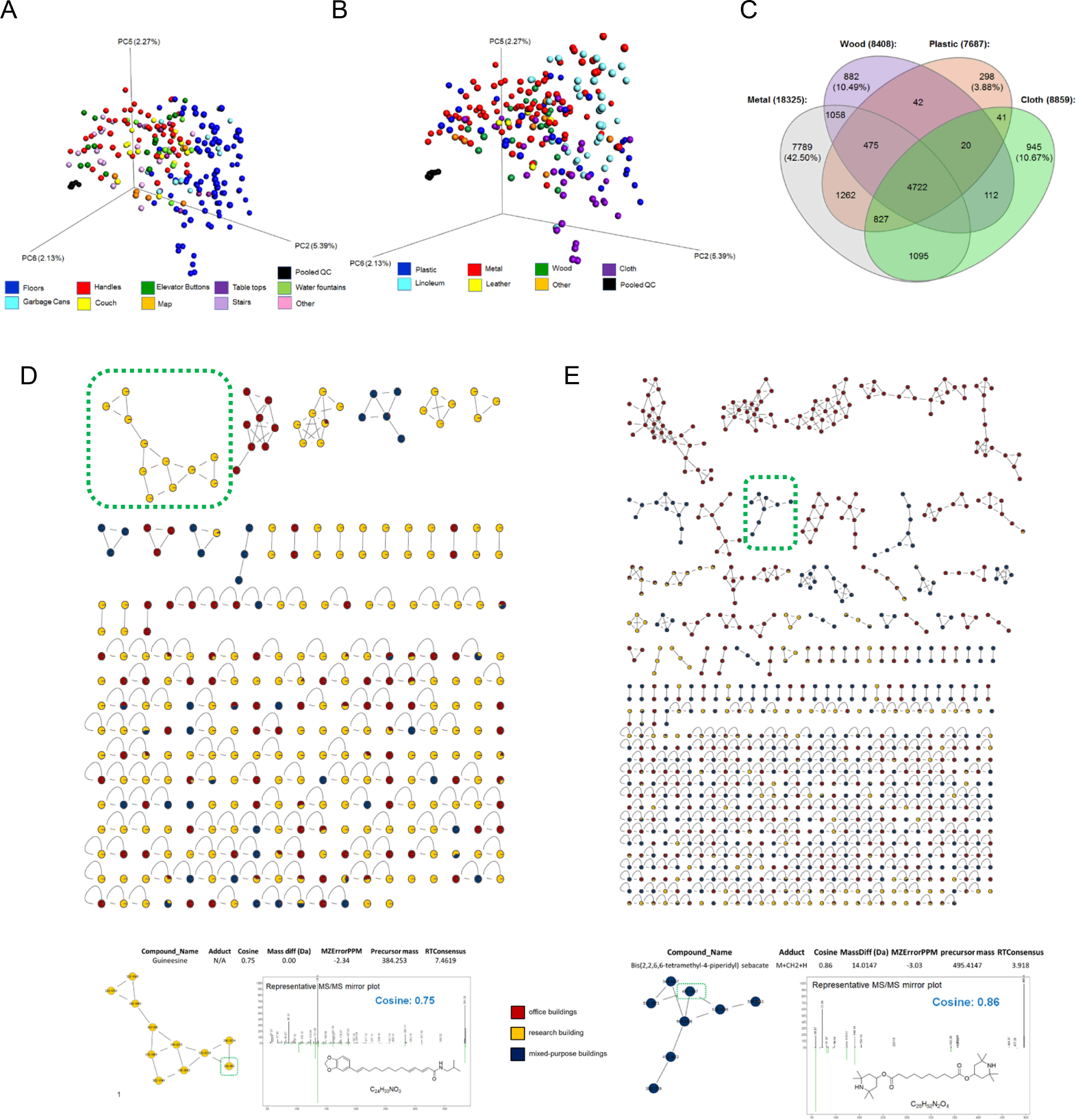
Significant impact of surface material on the recovered chemical profile. (A) Principal coordinate analysis showing segregation of samples based on sampling location within the three building types (Canberra distance metric; p<0.001 PERMANOVA). (B) Significant clustering of samples based on surface material, by principal coordinate analysis (Canberra distance metric; p<0.001 PERMANOVA). (C) Venn diagram showing the percent of recovered molecules unique to each material type. Metal surfaces yielded the highest numbers of unique chemical features. (D-E) Chemical families unique to each surface material. (D) Subnetworks of unique compounds identified from plastic (top) and representative chemical family (bottom). (E) Subnetworks of unique compounds identified from wood (top) and representative chemical family (bottom).

Different sampling locations were often made of different materials, so we also assessed the contribution of the surface material to the recovered chemical profile. For simplicity, the types of materials were filtered down to plastic, metal, cloth, wood, linoleum, and leather (**Fig. 2B**). Using the same Canberra distance matrix, we observed statistically significant clustering of samples based on the surface material (**Fig. 2B** PERMANOVA p<0.001). The greatest diversity of unique chemicals were recovered from metal surfaces (**Fig. 2C, Fig. S9**), which could be because most samples collected from metal surfaces are from commonly-touched places of research buildings or offices (door handles, elevator buttons etc.). Chemical families identified from metal surfaces include food constituents, personal care products and home use products (**Fig. S9**). The unique chemical families identified from the plastic surfaces mainly come from food or cleaning products (*e.g.* guineesine, a compound found in pepper; **Fig. 2D**), while the chemicals identified from the wood surface may come from coatings or personal care products (*e.g.* Bis(2,2,6,6-tetramethyl-4-piperidyl) sebacate; **Fig. 2E**). From the cloth surfaces, compounds from plants were often identified, possibly due to their location in building lunchrooms where food is likely to be spilled (**Fig. S8**).

Finally, we investigated features common across all buildings. These represent chemicals that most people are likely to encounter over the course of their workweek. Molecules commonly detected across all buildings include biological derivatives (fatty acids, amino acids, other related biomolecules), commonly found on high-touch surfaces which were not regularly cleaned (e.g. elevator buttons, door handles/knobs), detergents and molecules found in cleaning products (cocamidopropylbetaine derivatives, benzyldimethylstearylammonium cation derivatives) and natural product phytochemicals (presumably from food sources and personal care products). Compounds such as tangeritin (a natural compound found in the peels of tangerines and other citrus fruits) and piperine (a natural compound found in black pepper) were found on commonly touched surfaces, as well as surfaces used in the preparation and/or consumption of food products. Plasticizers, such as phthalate derivatives, were identified on multiple surfaces across all building types and on multiple materials. Finally, pharmaceutical derivatives such as levorphanol and clotrimazole were identified on multiple commonly touched surfaces across all building types and multiple surface types. Interestingly, both topical and orally ingested pharmaceuticals were identified on similar surfaces. This suggests that the scope of passive pharmaceutical exposure expands beyond topical formulations (**Table 3, Fig. S10**).

## Discussion

Overall, our results support shaping of the building surface metabolome by building function (**Fig. 1**). We identified distinct overall chemical profiles between research, office and mixed-purpose buildings, including many molecules that are likely occupant-derived (*e.g.* palmitoyl ethanolamide). Although our study sampled only building surfaces and as such was not designed to assess molecule sources, many of the molecules we detected were also found in other LC-MS/MS analyses of the human skin and frequently touched objects, further supporting a human source (over 8,000 matches with MassIVE datasets MSV000079181, MSV000078683, MSV000078622, MSV000080031, MSV000079558, MSV000078556, MSV000078816, MSV000078556, MSV000078816, MSV000078832, MSV000078993, MSV000079389, MSV000078556 [8][20,21] and **Table 2**). The ability to link our data to prior metabolomics studies therefore strongly enhanced this data’s usefulness. Future work will expand our study to investigate molecule transference from building surfaces to worker hands and vice versa.

Several food-derived molecules were found at higher levels in research buildings than in office buildings. This is likely due to the fact that the public areas sampled in the research buildings include the lunch room, while the office buildings do not have meal areas, and occupants either eat outside the building or at their desks (not sampled). Likewise, the prevalence of food molecules on cloth surfaces represent their presence on the chairs in these meal areas. The higher prevalence of palmitoyl ethanolamide and medications in the mixed-use building likely reflects its high-traffic nature, whereas fewer people frequent the research and office buildings. We also observed a significant impact of sampling site (location and material, **Fig. 2**) on the overall recovered metabolite profile. This latter observation highlights the importance of standardizing sampling locations, as implemented here. Our selection of five buildings within the same organization (with the office and research buildings in the same research park) also helped limit possible confounders due for example to differential cleaning practices across organizations.

Some of the detected molecules may present a health risk. Phthalates for example have been linked to asthma and allergies [22]; exposure to detergents such as cocamidopropylbetaine (*m/z* 301.285 RT 5.22 min, detected in all building types) can cause allergic contact dermatitis [23]. However, it is important to note that only 34% of our dataset had family-level annotations, with 2.5% of molecules receiving compound-level annotations (level 2 confidence per metabolomics standards initiative [17]). This highlights the major challenge of metabolomics studies of human-building interactions, the un-annotated “dark matter” [24]. Linking molecules detected in one study with other LC-MS/MS studies of the built environment can help shed at least some insight on these molecules. Indeed, our results integrate well with prior studies of the built environment, with 21,185 matches to molecules in other studies of the built environment (out of 127,397 total dataset matches; MassIVE accession numbers MSV000079720, MSV000079714, MSV000079717, MSV000079706, MSV000079709 [7]). Annotated molecules shared between our study and this prior work include detergents (*e.g.* cocamidopropylbetaine), food products (*e.g.* constiuents of pepper), illicit drugs (cocaine) and medications (*e.g.* erythromycin). Although the presence of cocaine in these settings may seem surprising, it is commonly found on US currency [25] and was previously reported in other studies of the built environment [7].

In conclusion, our results highlight the applicability of LC-MS/MS to study building-occupant interactions and to identify workplace chemical exposure risks, in a targeted setting. Future work will be required to assess whether detected molecules present a health risk to employees and building visitors.

## Acknowledgements

The authors would like to thank all the building managers who allowed us to collect samples from their buildings. The authors would also like to thank Wen Yang for helping with sample collection, and Shelley Kane and Adwaita Parab for assistance with molecule extraction. This work was supported by start-up funds from the University of Oklahoma.

## Abbreviations

LC-MS/MS: liquid chromatography-tandem mass spectrometry
m/z: mass over charge ratio
RT: retention time

