## Supplemental data for "Characterization of the workplace chemical exposome using untargeted LC-MS/MS: a case study"

**Figure S1. Euler diagram showing significant overlap between all buildings.**

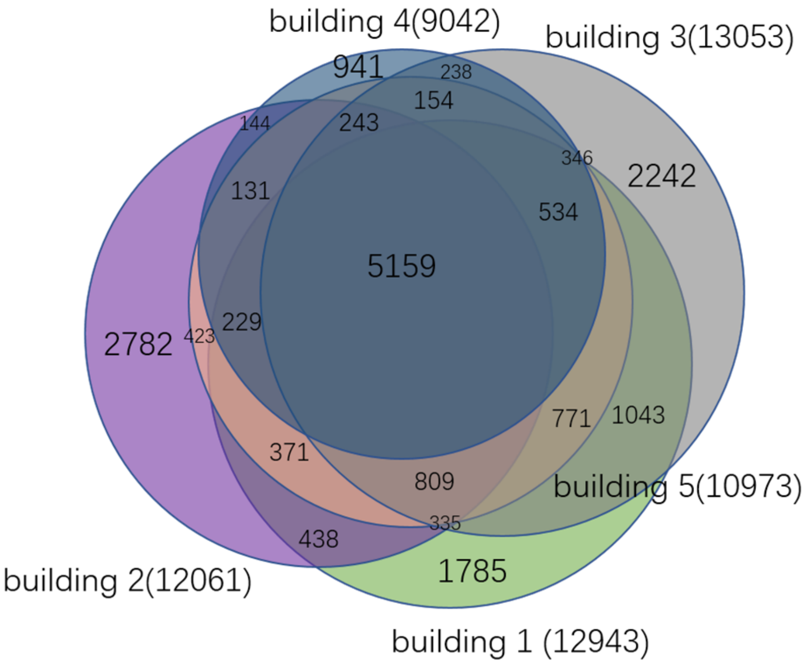

**Figure S2: GNPS mirror plots for molecules differing between building types.** Top, experimental MS2 spectrum (black). Bottom, reference library spectrum (green). (A) *m/z* 223.095 RT 4.72 min match to diethyl phthalate. (B) *m/z* 279.232 RT 7.06 min match to methyl gamma-linolenate. (C) *m/z* 286.143 RT 5.21 min match to piperine. (D) *m/z* 286.274 RT 4.85 min match to N-(2-hydroxypropyl)dodecanamide. (E) *m/z* 299.162 RT 4.05 min match to 1,7-bis(4-hydroxyphenyl)heptane-3,5-diol. (F) *m/z* 300.289 RT 7.42 min match to palmitoyl ethanolamide. (G) *m/z* 343.188 RT 4.94 min match to tris(2-butoxyethyl) phosphate (-$C_4H_8$ to [M+H]+ adduct). (H) *m/z* 365.17 RT 4.94 min match to tris(2-butoxyethyl) phosphate (-$C_4H_8$ to [M+Na]+ adduct). (I) *m/z* 387.200 RT 4.34 min match to (4S)-4-hydroxy-3,5,5-trimethyl-4-[(E)-3-[(2R,3R,4S,5S,6R)-3,4,5-trihydroxy-6-(hydroxymethyl)oxan-2-yl]oxybut-1-enyl]cyclohex-2-en-1-one. (J) *m/z* 421.232 RT 6.74 min match to tris(2-butoxyethyl) phosphate ([M+Na]+). (K) *m/z* 819.474 RT 6.74 min match to tris(2-butoxyethyl) phosphate ([2M+Na]+).0

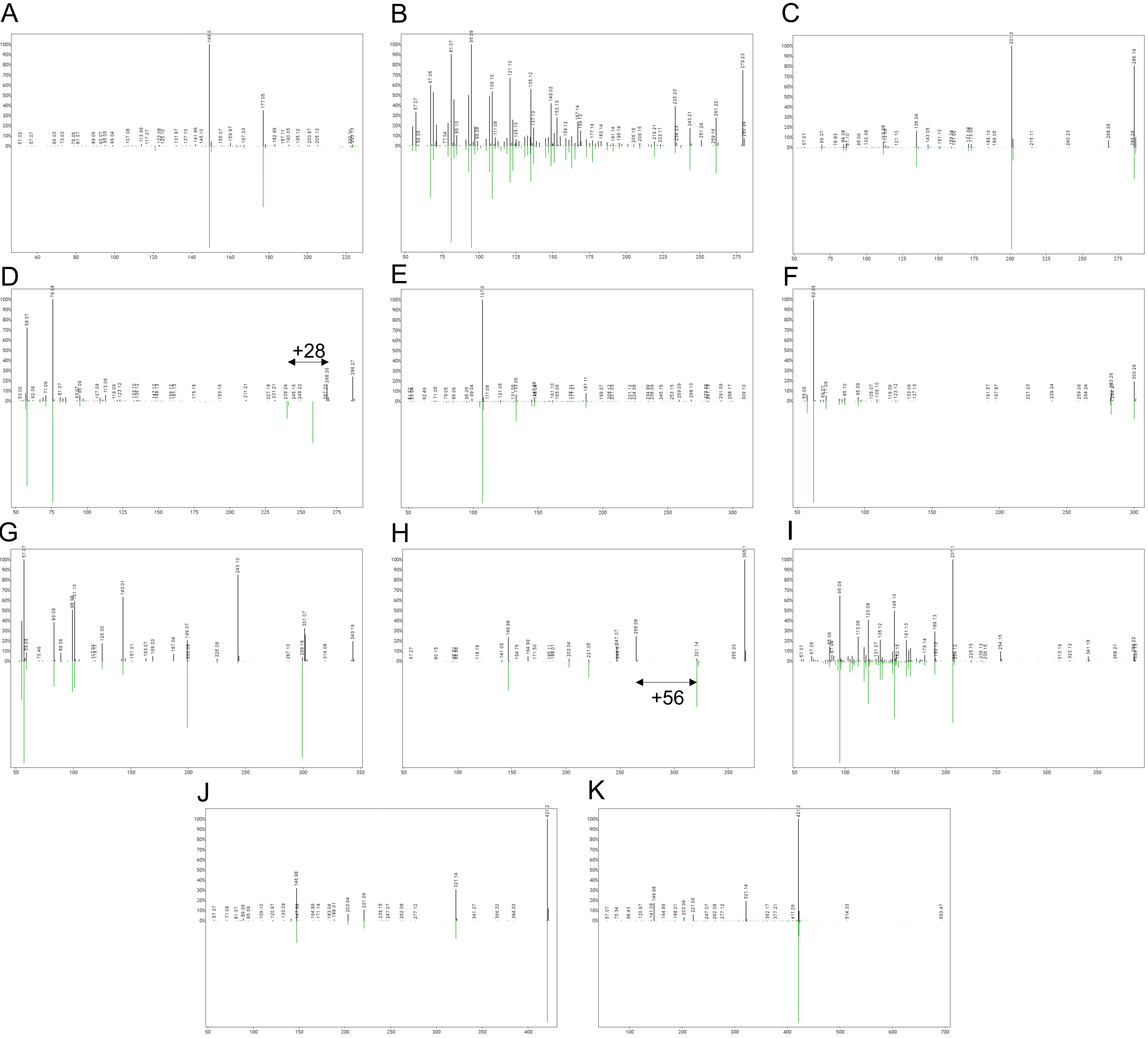

**Figure S3. Principal coordinate analysis by building.** Buildings of a given type do not segregate from each other.

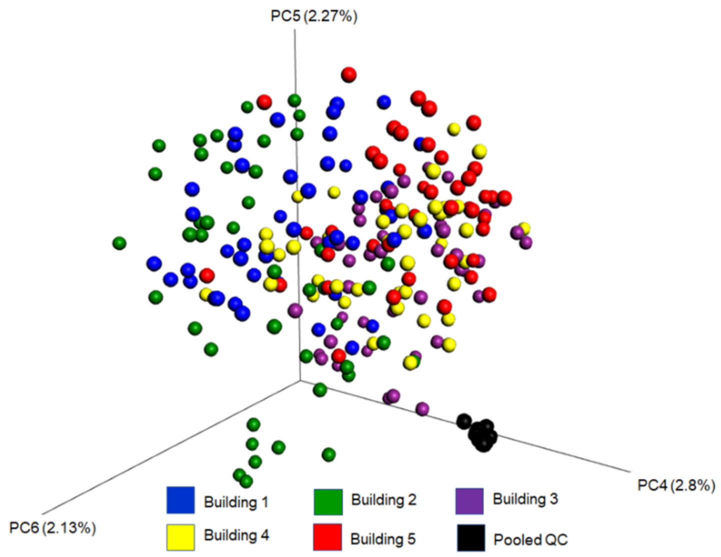

**Figure S4: Differential abundance of features differing by building type (as identified by random forest).** (A) *m/z* 115.075 RT 6.39 min. (B) *m/z* 121.04 RT 0.37 min. (C) *m/z* 139.05 RT 0.31 min. (D) *m/z* 149.023 RT 4.72 min. (E) *m/z* 223.095 RT 4.72 min. (F) *m/z* 272.258 RT 4.75 min. (G) *m/z* 279.232 RT 7.06 min. (H) *m/z* 286.143 RT 5.21 min. (I) *m/z* 286.274 RT 4.85 min. (J) *m/z* 299.162 RT 4.05 min. (K) *m/z* 300.289 RT 7.42 min. (L) *m/z* 343.188 RT 4.94 min. (M) *m/z* 343.211 RT 5.18 min. (N) *m/z* 365.17 RT 4.94 min. (O) *m/z* 386.238 RT 2.46 min. (P) *m/z* 387.2 RT 4.34 min. (Q) *m/z* 413.229 RT 6.39 min. (R) *m/z* 421.232 RT 6.74 min. (S) *m/z* 425.252 RT 3.91 min. (T) *m/z* 435.211 RT 6.39 min. (U) *m/z* 470.369 RT 6.12 min. (V) *m/z* 504.319 RT 3.95 min. (W) *m/z* 509.273 RT 3.95 min. (X) *m/z* 514.395 RT 6.13 min. (Y) *m/z* 548.345 RT 4.02 min. (Z) *m/z* 553.299 RT 4.02 min. (AA) *m/z* 592.37 RT 4.08 min. (AB) *m/z* 597.324 RT 4.07 min. (AC) *m/z* 602.446 RT 6.14 min. (AD) *m/z* 819.474 RT 6.74 min.

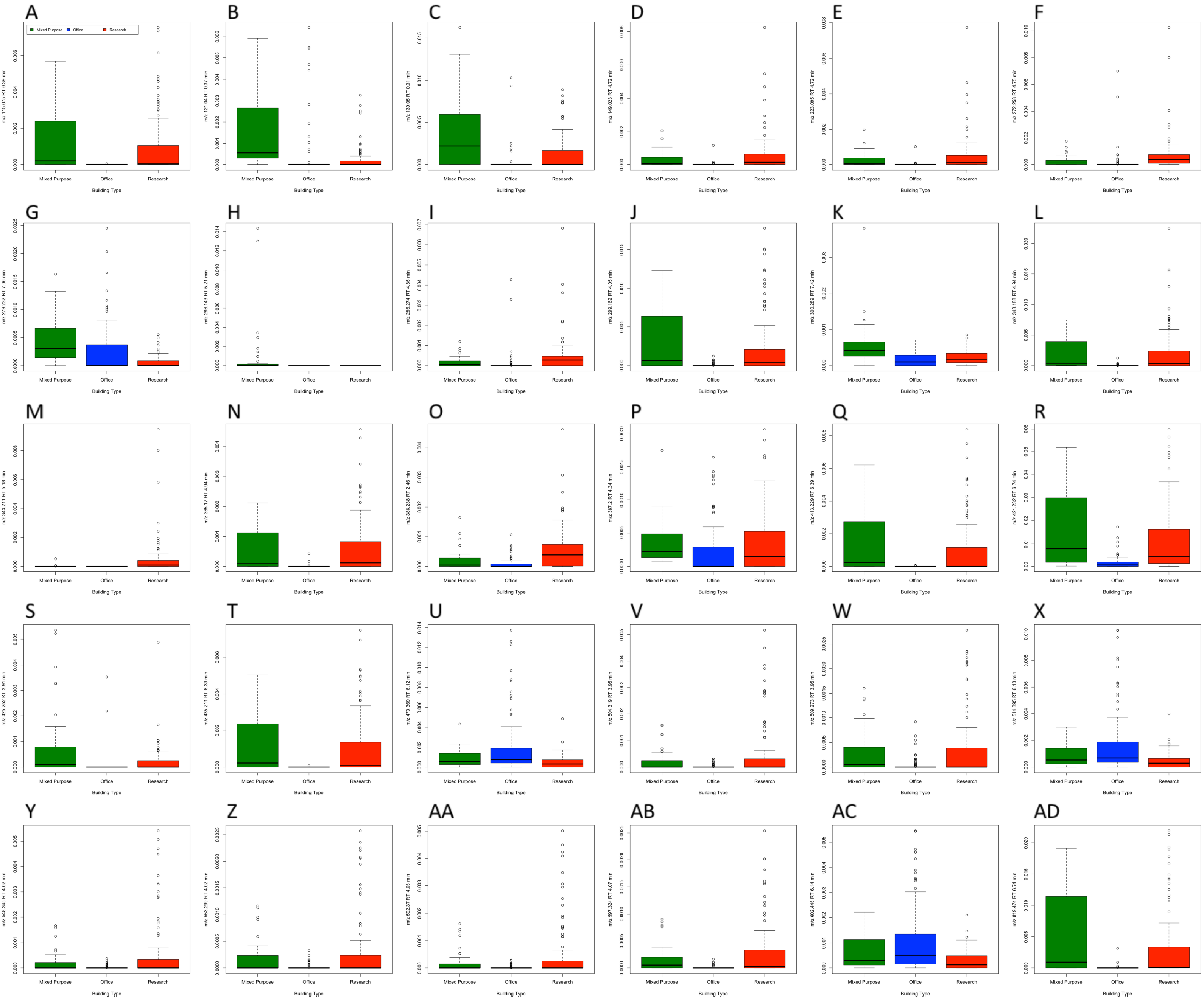

**Figure S5. Per-building abundance of top 30 most differential features, as identified by random forest and with statistically significant ANOVA results (*p*-value < 0.05), including:** (A) *m/z* 115.075 RT 6.39 min, (B) *m/z* 121.04 RT 0.37 min, (C) *m/z* 139.05 RT 0.31 min, (D) *m/z* 149.023 RT 4.72 min, (E) *m/z* 223.095 RT 4.72 min (F) *m/z* 272.258 RT 4.75 min, (G) *m/z* 279.232 RT 7.06, (H) *m/z* 286.143 RT 5.21 min, (I) *m/z* 286.274 RT 4.85 min, (J) *m/z* 299.162 RT 4.05 min, (K) *m/z* 300.289 RT 7.42 min, (L) *m/z* 343.188 RT 4.94 min, (M) *m/z* 343.211 RT 5.18 min, (N) *m/z* 365.17 RT 4.94 min, (O) *m/z* 386.238 RT 2.46 min, (P) *m/z* 413.229 RT 6.39 min, (Q) *m/z* 421.232 RT 6.74 min, (R) *m/z* 425.252 RT 3.91 min, (S) *m/z* 435.211 RT 6.39 min, (T) *m/z* 470.369 RT 6.21 min, (U) *m/z* 504.319 RT 3.95 min, (V) *m/z* 509.273 RT 3.95 min, (W) *m/z* 514.395 RT 6.13 min, (X) *m/z* 548.345 RT 4.02 min, (Y) *m/z* 553.299 RT 4.02 min, (Z) *m/z* 592.37 RT 4.08 min, (AA) *m/z* 597.324 RT 4.07 min, (AB) *m/z* 602.446 RT 6.14 min, and (AC) *m/z* 819.474 RT 6.74 min.

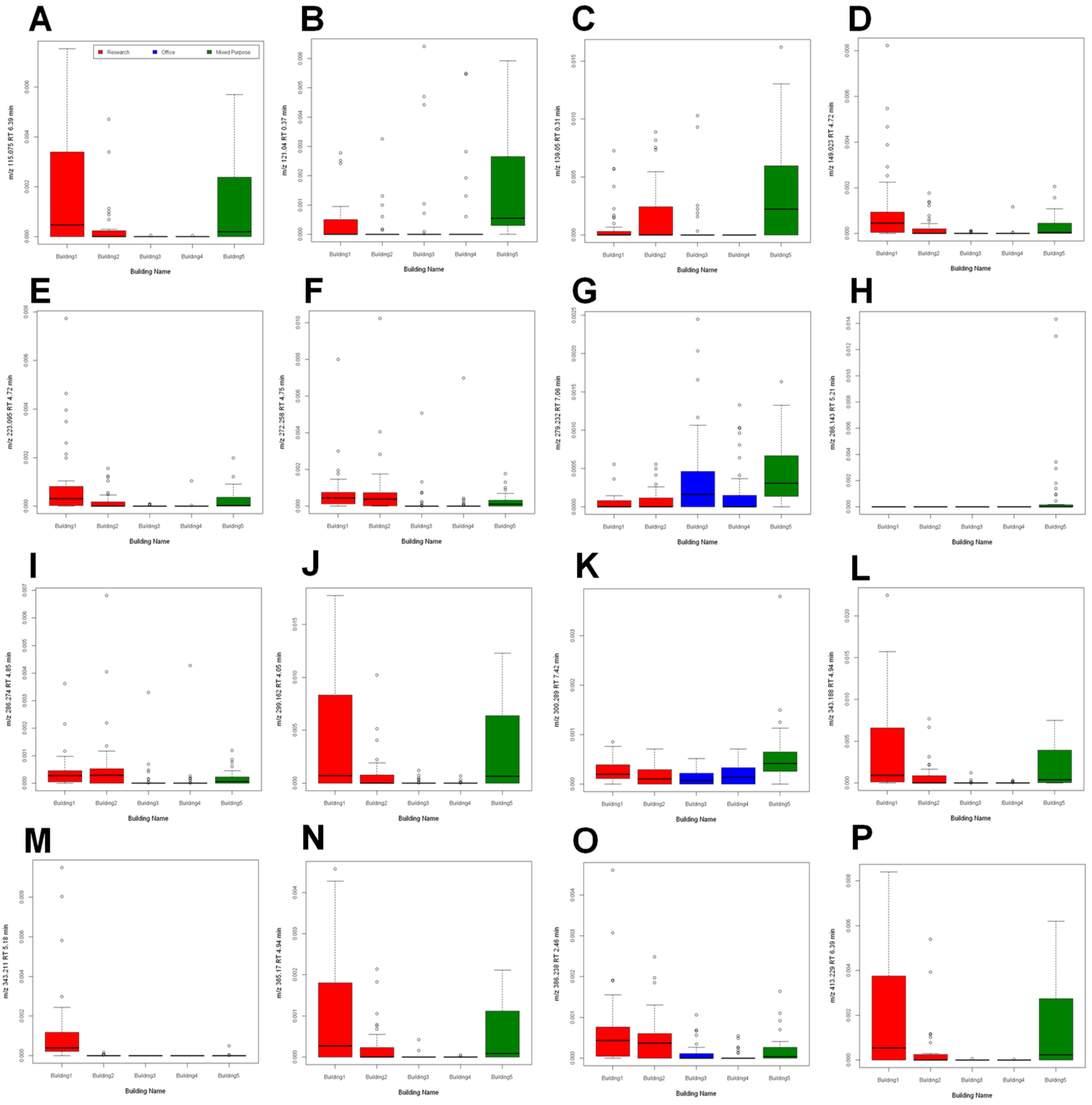

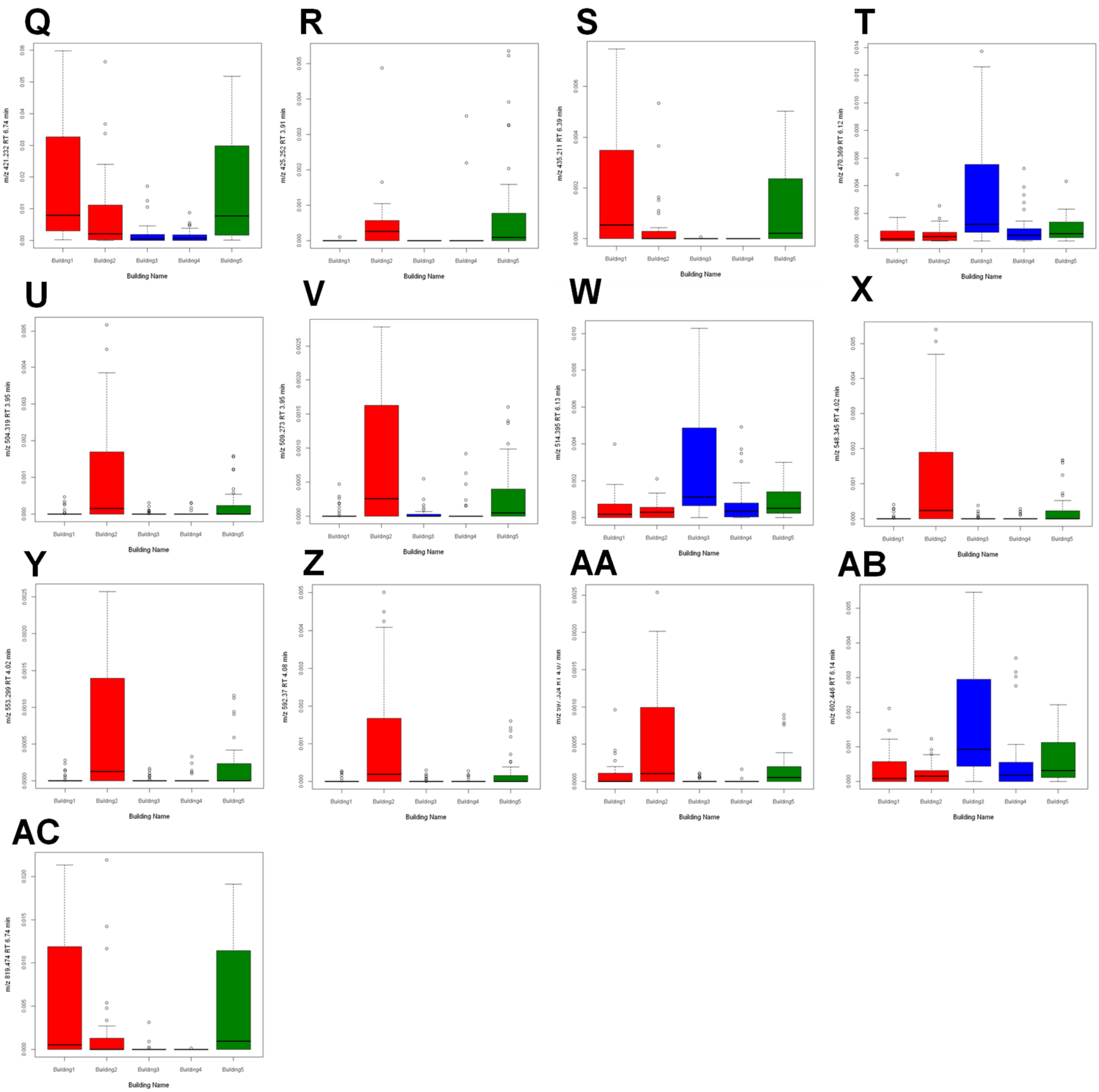
Q
R
S
T
U
V
W
X
Y
Z
AA
AB
AC
m/z 421.232 RT 6.74 min
m/z 425.252 RT 3.91 min
m/z 435.211 RT 6.39 min
m/z 470.369 RT 6.12 min
m/z 504.319 RT 3.95 min
m/z 509.273 RT 3.95 min
m/z 514.395 RT 6.13 min
m/z 548.345 RT 4.02 min
m/z 553.299 RT 4.02 min
m/z 592.37 RT 4.08 min
m/z 602.446 RT 6.14 min
m/z 819.474 RT 6.74 min
Building1
Building2
Building3
Building4
Building5
Building Name

**Figure S6. Overall molecular network.**

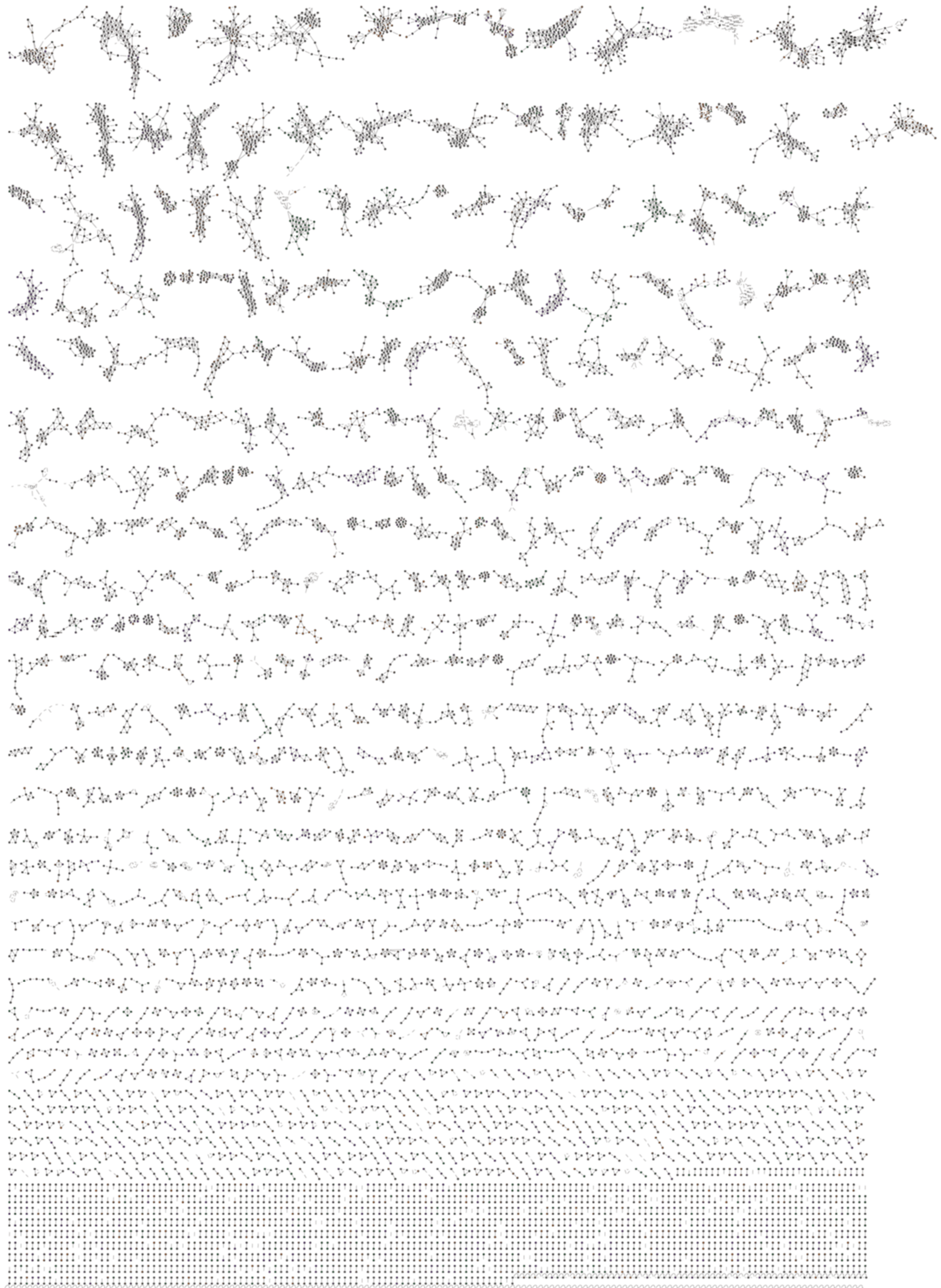

**Figure S7. Additional representative compounds identified from wood-specific subnetworks:** plasticizer bis(2-ethylhexyl) phthalate.

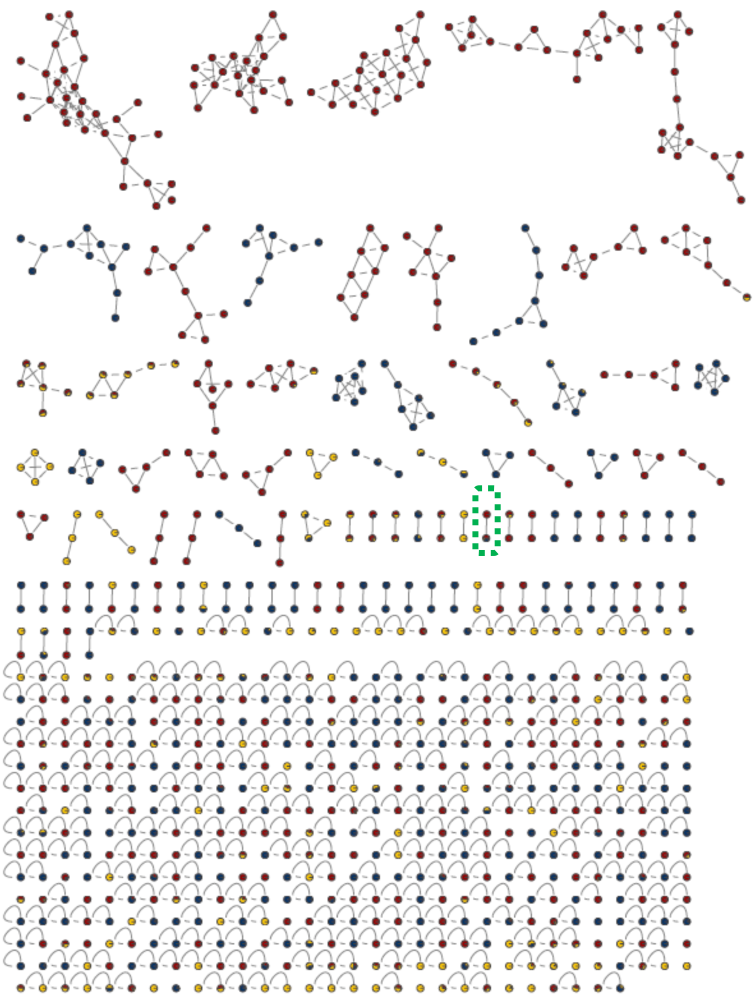

| Compound_Name | Adduct | Cosine | MassDiff (Da) | MZErrorPPM | Precursor mass | RTConsensus |
|---|---|---|---|---|---|---|
| Bis(2-ethylhexyl) phthalate | M+H | 0.98 | 0.00 | 0.467961 | 391.2838 | 1.7649 |
| Bis(2-ethylhexyl) phthalate | M+H | 0.98 | 0.00 | 0.779934 | 391.2837 | 2.7952 |

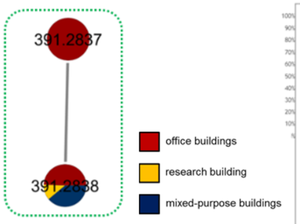


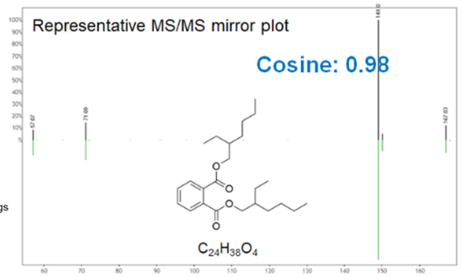

**Figure S8. Additional representative compounds identified from cloth-specific subnetworks.** Identifiable molecules include kaempferol-3-glucuronide, a plant- and fruit flavonoid, and gamma-dodecalactone, a fruit lactone.

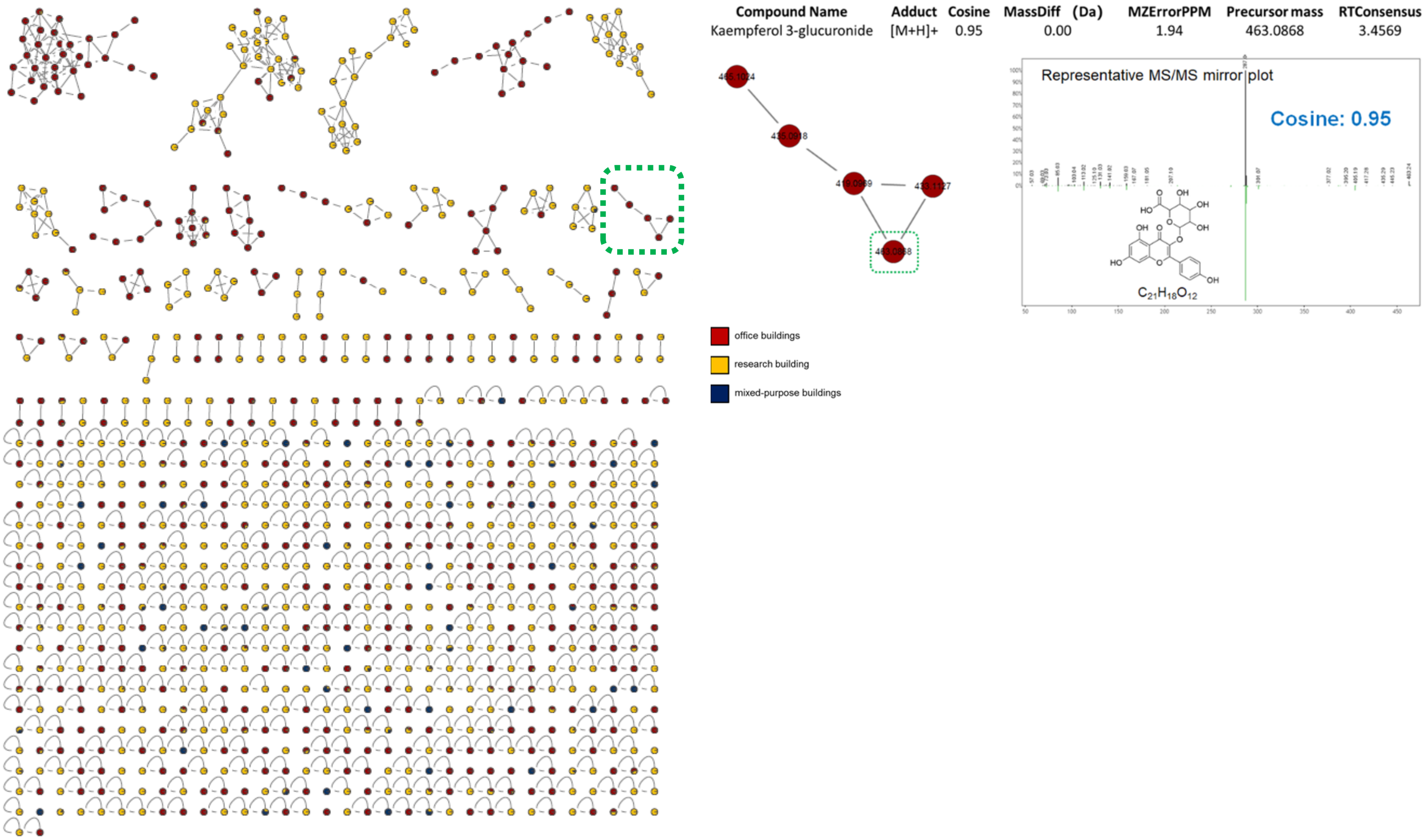

**Figure S9. Additional representative compounds identified from metal-specific subnetworks.** Identifiable molecules include benzyl nicotinate (skin care product); gamma-dodecalactone, a fruit lactone; tetraethylene glycol (cleaning product).

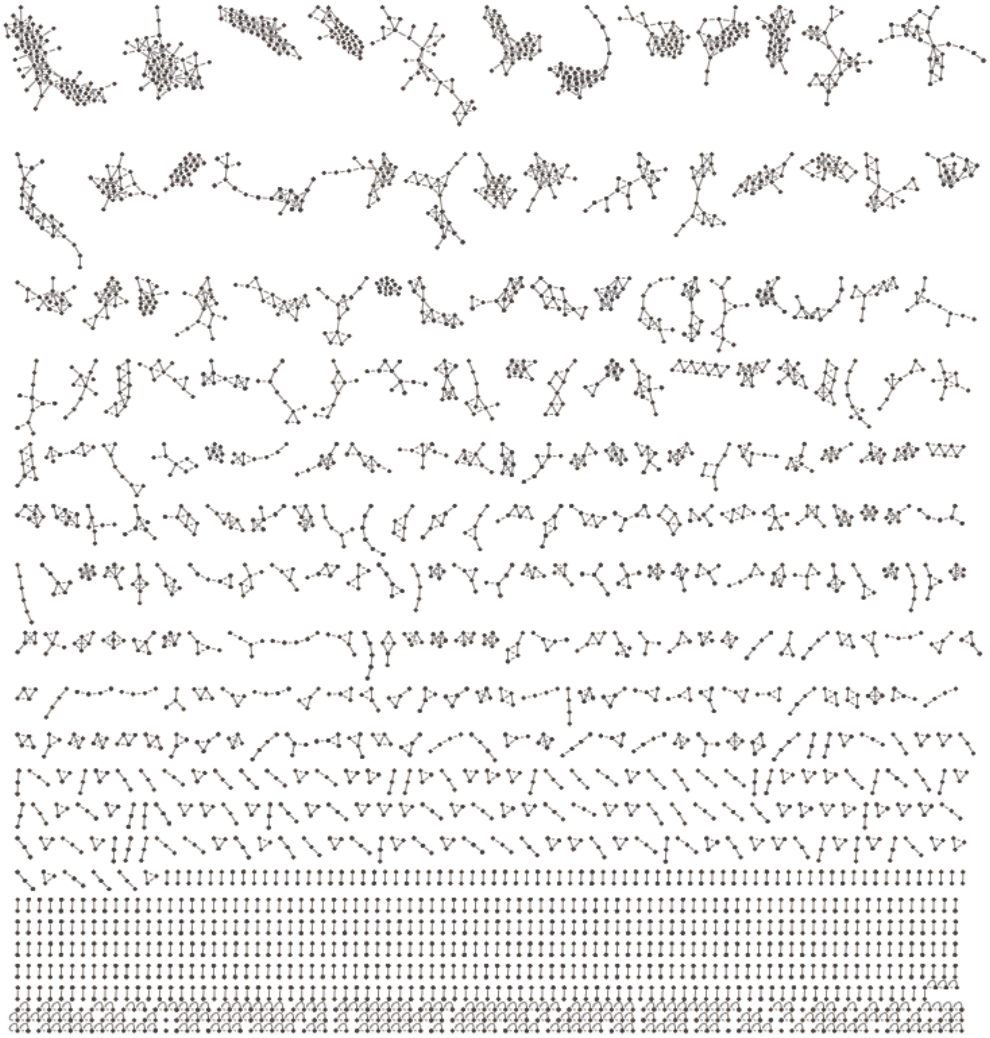

| Compound Name | Adduct | Cosine | MassDiff | MZErrorPPM | Precursor mass | RTConsensus |
|---|---|---|---|---|---|---|
| Benzyl nicotinate | M+H | 0.91 | 0.00 | -3.27 | 214.0861 | 4.6627 |

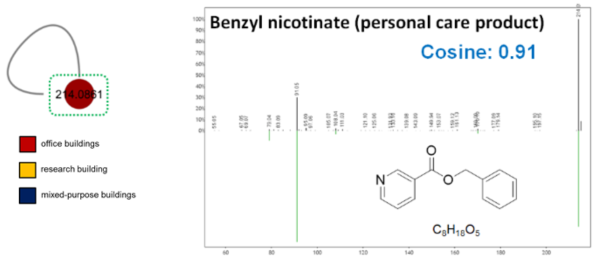


| Compound Name | Adduct | Cosine | Mass difference | MZErrorPPM | Precursor mass | RTConsensus |
|---|---|---|---|---|---|---|
| gamma-Dodecalactone | M+H | 0.94 | 0.00 | -3.01 | 199.1692 | 5.5087 |

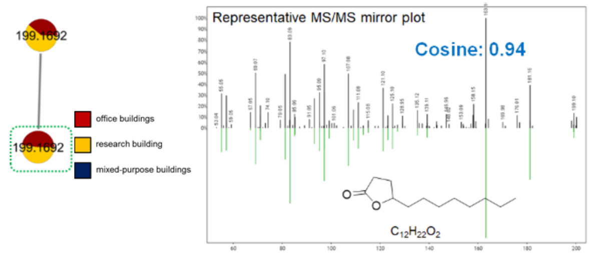


| Compound Name | Adduct | Cosine | MassDiff | MZErrorPPM | Precursor mass | RTConsensus |
|---|---|---|---|---|---|---|
| Tetraethylene glycol | M+H | 0.97 | 0.00 | -3.07 | 195.1226 | 7.2173 |

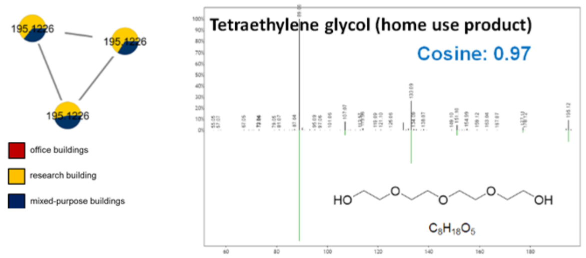

**Figure S10. GNPS mirror plots for molecules in Table 3.** Top, experimental MS2 spectrum (black). Bottom, reference library spectrum (green). (A) *m/z* 201.054 RT 5.48 min match to piperlongumine. (B) *m/z* 214.086 RT 4.66 min match to benzyl nicotinate. (C) *m/z* 239.106 RT 0.28 min match to 4-(2-hydroxyethyl)piperazine-1-ethanesulfonic acid (HEPES). (D) *m/z* 276.175 RT 4.46 min match to cyclobenzaprine. (E) *m/z* 734.468 RT 4.22 min match to erythromycin. (F) *m/z* 133.0637 RT 3.61 min match to levorphanol. (G) *m/z* 135.117 RT 4.85 min match to patchouli alcohol. (H) *m/z* 135.117 RT 6.62 min match to undecanedioic Acid. (I) *m/z* 205.097 RT 0.87 min match to tryptophan. (J) *m/z* 219.174 RT 4.41 min match to caryophyllene oxide. (K) *m/z* 219.174 RT 6.22 min match to nootkatone. (L) *m/z* 237.221 RT 6.32 min match to palmitelaidic acid. (M) *m/z* 277.0776 RT 4.79 min match to clotrimazole. (N) *m/z* 286.1434 RT 5.49 min match to piperine. (O) *m/z* 301.285 RT 5.22 min match to cocamidopropyl betaine. (P) *m/z* 304.154 RT 3.27 min match to cocaine. (Q) *m/z* 356.243 RT 6.99 min match to piperonyl butoxide. (R) *m/z* 360.362 RT 7.68 min match to benzyldimethylstearylammonium cation. (S) *m/z* 369.351 RT 10.49 min match to cholesterol. (T) *m/z* 373.1275 RT 5.49 min match to tangeritin. (U) *m/z* 391.284 RT 6.83 min match to dioctyl phthalate. (V) *m/z* 415.254 RT 7.51 min match to nonaethylene glycol. (W) *m/z* 439.356 RT 4.70 min match to Oleanolic Acid. (X) *m/z* 465.103 RT 3.25 min match to astragalin (+O derivative). (Y) *m/z* 565.118 RT 3.66 min match to Quercetin 3-O-malonylglucoside (+$CH_2$ derivative). (Z) *m/z* 757.217 RT 3.02 min match to 5,7-dihydroxy-2-[4-[(2S,3R,4S,5S,6R)-3,4,5-trihydroxy-6-(hydroxymethyl)oxan-2-yl]oxyphenyl]-3-[(2S,3R,4S,5S,6R)-3,4,5-trihydroxy-6-[[(2R,3R,4R,5R,6S)-3,4,5-trihydroxy-6-methyloxan-2-yl]oxymethyl]oxan-2-yl]oxychromen-4-one.

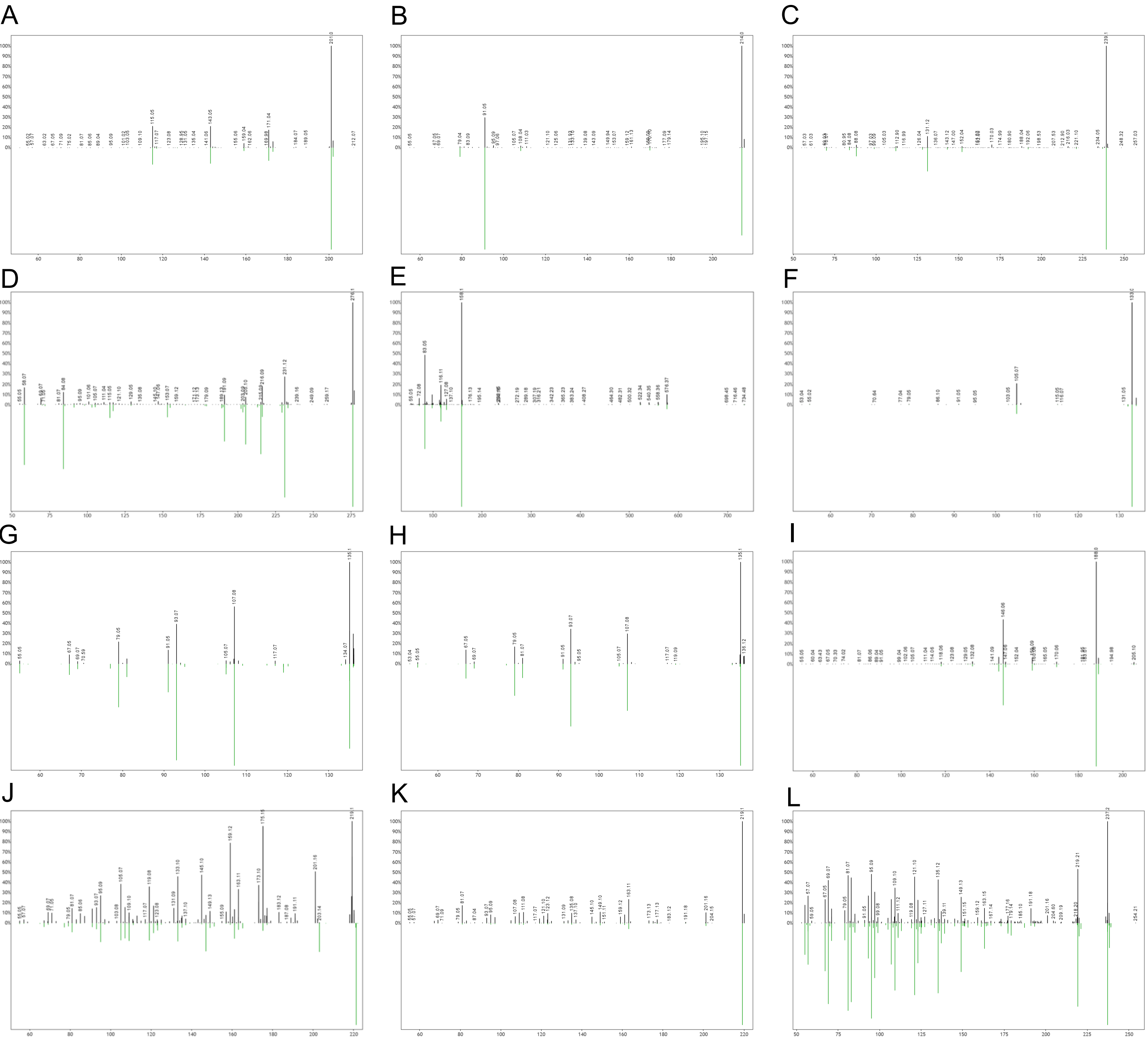

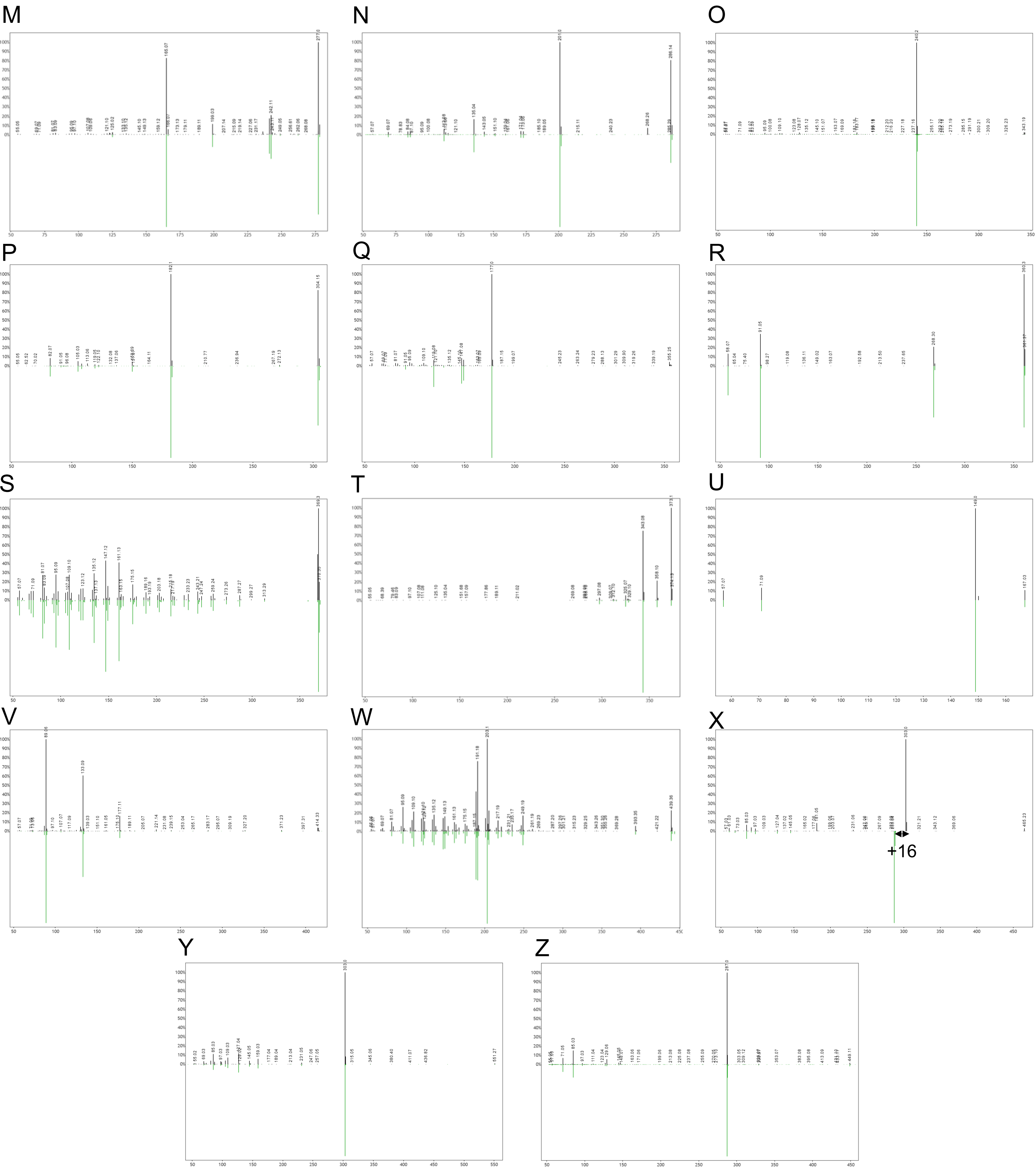
M
N
O
P
Q
R
S
T
U
V
W
X
+16
Y
Z
